# Application of barcode sequencing to increase the throughput and complexity of *Plasmodium falciparum* genetic screening

**DOI:** 10.1101/2024.09.05.611197

**Authors:** Allan Muhwezi, Mehdi Ghorbal, Theo Sanderson, Maria Ivanova, Rizwan Ansari, Sarah Harper, Wesley Wong, Reiner Schulte, Gareth Girling, Frank Schwach, Ellen Bushell, Charlotte Beaver, Oliver Billker, Julian C. Rayner

**Affiliations:** Cambridge Institute for Medical Research, University of Cambridge, Cambridge, UK; Wellcome Sanger Institute, Wellcome Genome Campus, Cambridge, UK; Harvard T.H. Chan School of Public Health, Harvard University, Cambridge, USA; Umeå University, Umeå, Sweden

**Keywords:** *Plasmodium falciparum*, malaria, CRISPR-Cas9, experimental genetics, pooled phenotyping, parasite growth, DNA replication, cell cycle

## Abstract

All the pathology and symptoms associated with malaria are caused by the growth of *Plasmodium* parasites inside human red blood cells. This process, which in the case of the major human malaria pathogen *Plasmodium falciparum* takes place over a 48-hour period, involves multiple tightly regulated developmental transitions. Understanding the *P. falciparum* genes that regulate these key processes could lead to the identification of targets for new drugs. However, while large-scale sequencing efforts have led to a good understanding of the *P. falciparum* genome and how it evolves over time and space, a disconnect remains between the amount of genome sequence data available and the amount of data describing what exactly the genes contained within it do – the phenotype. We have generated a panel of 66 *P. falciparum* lines carrying individual gene knockouts tagged with unique DNA barcodes. We then used these lines in a series of assays that combine flow cytometry, cell sorting and DNA barcode quantification using next generation sequencing (Barcode Sequencing or BarSeq) to phenotype key aspects of the parasite life cycle such as growth, replication capacity and cell cycle progression. This approach both yields new data about individual gene function, and outlines a new approach where barcoded *P. falciparum* lines are used in pooled BarSeq-based assays to generate more precise phenotype data at scale.

## Introduction

*Plasmodium falciparum* parasites cause the most virulent form of human malaria which accounts for hundreds of millions of cases and more than 500,000 deaths annually (WHO 2022). Significant progress has been recently made in controlling this disease, but the repeated emergence and spread of antimalarial drug and insecticide resistance, along with currently limited vaccine coverage, all suggest that further interventions will be required to deliver malaria elimination (Donnelly, Isaacs and Weetman 2016, WHO 2022). The search for new antimalarial targets can be directly informed by the *P. falciparum* genome, which encodes over 5400 genes (Gardner, Hall et al. 2002). Homology and functional prediction reveals many potential intervention targets, including multiple enzymes and transporters, but more than 50% of *P. falciparum* genes lack conservation with model organisms and therefore their function could not be directly predicted by homology alone (Gardner, Hall et al. 2002). Despite extensive annotation efforts, the function of ∼35% of the *P. falciparum* genome remain hypothetical (Oberstaller, Otto et al. 2021). More targeted and, critically, scalable efforts are needed to understand *P. falciparum* gene function in order to identify new drug and vaccine targets.

Experimental genetic manipulation of the *Plasmodium* genome has been central to efforts to characterize gene function since it was first established in the early 1990’s (Van Dijk, Waters and Janse 1995, Wu, Sifri et al. 1995). In the subsequent decades, genetics has been used to identify genes involved in many parasite processes such as erythrocyte invasion, drug resistance, metabolism, and virulence. This has been achieved by targeting DNA loci with a range of manipulation strategies such as knock-out, replacement or mutagenesis using either conventional recombination approaches or targeted recombination using CRISPR/Cas9 and zinc finger nucleases (Straimer, Lee et al. 2012, Ghorbal, Gorman et al. 2014). A range of complex conditional tools have also been developed to study the function of essential genes/proteins, including mRNA degradation using inducible promoters and *glm S* ribozymes, protein degradation using destabilizing domains such as DD and DDD (de Koning-Ward, Gilson and Crabb 2015) and conditional gene deletion using small-molecule induced diCre and FRT recombinases (Combe, Giovannini et al. 2009, Collins, Das et al. 2013). These detailed gene-by-gene approaches have been invaluable and have produced a wealth of insight and data. However many of these approaches are relatively low throughput due to low transfection efficiencies, limited numbers of selectable markers and the need to generate complex vectors and constructs (de Koning-Ward, Gilson and Crabb 2015, Lee, Lindner et al. 2019). More systematic high throughput approaches are needed to both the generation and phenotyping of *P. falciparum* mutant lines to truly expand understanding of the many genes of unknown function.

Despite these challenges, there have been some large genetic screens in *Plasmodium* parasites. Before the development of CRISPR/Cas9, two medium-scale screens of >50 genes encoding kinases or proteins predicted to be involved in cytoadherence were carried out with standard recombination technology (Maier, Rug et al. 2008, Solyakov, Halbert et al. 2011). For the rodent model *P. berghei,* the *Plasmo*GEM (*Plasmodium* Genetic Modification) Project used a lambda recombination and Gateway cloning system to generate targeted barcoded vectors for >2500 genes, which were transfected in pools of 100 vectors at a time. Nextgen sequencing was then used to quantify the relative abundance of each barcode within the mixed pool (Barcode Sequencing or BarSeq) in order to quantify individual parasite growth rates (Bushell, Gomes et al. 2017). This approach has now been used successfully to assign growth phenotypes at blood (Bushell, Gomes et al. 2017), pre-erythrocytic and mosquito (Stanway, Bushell et al. 2019) and gametocyte (Russell, Sanderson et al. 2023) stages in *P. berghei*. The lower transfection efficiency in *P. falciparum* has to date precluded a similar approach in the human pathogen, but here the *piggyBac* transposon system, which uses random insertion of *the piggyBac* transposon at TTAA sites, which occur at high frequencies in the AT-rich *P. falciparum* genome, has been used to generate >30,000 *P. falciparum* mutants and provide a first genome-wide estimate of gene essentiality in this species, based on the pattern of transposon insertion (Zhang, Wang et al. 2018). Both these large-scale screens show that a significant proportion of the *Plasmodium* genome is required for blood-stage growth.

A major challenge however that still remains is the development of more complex screens that extend beyond just parasite growth measurements. Parasite growth could be impaired for multiple reasons, from limiting one particular step in intra-erythrocytic development (e.g., metabolism or invasion) or by extending the length of the intra-erythrocytic cycle as whole. In addition, the environment and growth conditions are also likely to influence gene function and phenotypes, as has been shown extensively in other organisms such as yeast (Hillenmeyer, Fung et al. 2008). Growth screens alone are therefore only the starting point for understanding gene function. In this work we explored the use of BarSeq to develop more systematic screening approaches specifically for *P. falciparum.* We did so by generating a library of 66 barcoded *P. falciparum* knockout lines using a custom CRISPR/Cas9 construct generation and transfection pipeline, and then screening them both as individual lines and as pools for parasite growth, DNA replication, intra-erythrocytic cycle length and progression, leveraging BarSeq as a method of quantification wherever possible. This generated both information about individual gene function, and also the potential for future larger-scale targeted genetic screens in *P. falciparum* which will accelerate the discovery of new potent drugs and combinatorial therapies with novel cellular targets through screening diverse libraries of small molecules as well as pinpointing gene function across multiple parasite lifecycle stages and environmental perturbations.

## Results

### Generation of a panel of *Plasmodium falciparum* deletion lines using a CRISPR/Cas9 pipeline

We developed a CRISPR/Cas9 pipeline to generate a panel of barcoded *P. falciparum* deletion lines for use in mixed pool phenotyping (**Supplementary Figure 1**). Guide RNA (gRNA) and homology repair designs were generated targeting 522 genes using an automated *in silico* construct design pipeline (see Methods). The homologous repair plasmid consists of a positive expression cassette (hDHFR, which can be selected using WR99210) flanked by 5’ and 3’ regions designed to target repair of the gRNA cleaved locus. An 11-12 nucleotide unique barcode flanked by conserved primer annealing sites was also incorporated into each homologous repair construct to facilitate subsequent high throughput phenotyping using barcode sequencing, BarSeq (Bushell, Gomes et al. 2017). In order to increase the chances of creating deletion lines with growth phenotypes, 392 (75.1%) genes were selected on the basis that deletion of their orthologues in *P. berghei* resulted in a slow growth phenotype (Bushell, Gomes et al. 2017). 31 genes predicted essential and 34 genes predicted dispensable in the same *P. berghei Plasmo*GEM screen were included as controls. An additional 65 genes predicted to be involved in invasion in *P. falciparum* and for which no clear orthologue could be identified in *P. berghei* were also included, again to increase the chance of revealing growth phenotypes.

Of the 522 construct designs, 234 (44.8%) were successfully cloned using a Gibson assembly pipeline and transfected by electroporation into *P. falciparum* parasite cultures enriched for ring stages. 152 (65.0%) transfectants were recovered after positive selection using the drug WR99210 and thereafter frozen down. Frozen isolates were subsequently revived, and a second round of positive selection performed, of which 93 (61.2%) lines regrew in the presence of WR99210. Positive/negative selection was then carried out to select against maintenance of the repair plasmid and hence for lines in which the homology region had integrated into the genome via double cross-over recombination; this resulted in the recovery of 66 (71.0%) unique lines. Once the combination of both positive and negative selection had been completed for all the lines, the nucleotide sequence identity of the barcode module was then determined using capillary sequencing, followed by PCR-based genotyping using primers flanking the integrated region in order to confirm cassette integration (see Methods). A summary of the proportion of genes, along with the estimated growth phenotypes based on the *Plasmo*GEM *P. berghei* screen (essential/slow/redundant) that progressed through each stage of the CRISPR/Cas9 pipeline is shown in **Table 1** (where the denominator used to calculate the % is the number of constructs/lines that passed the previous step). Details of each construct and its status along the transfection pipeline are included in **Supplementary Table 1**. In certain instances, >1 round of transfections was performed using the same constructs as shown in Supplementary Table 1. A summary of the genotyping results is shown in **Supplementary Table 2**. In general, there was some degree of concordance between predictions based on the *P. berghei Plasmo*GEM data and transfection outcome – of constructs that were transfected, only 2/14 genes (14.2%) identified as essential in *P. berghei* passed combined positive/negative selection, whereas 51/173 (29.5%) and 5/15 (33.3%) of genes identified as slow or redundant in *P. berghei* reached the same stage respectively.

**Table 1.** Outcome of CRISPR/Cas9 pipeline to generate barcoded gene deletion lines in *Plasmodium falciparum*.

| *Plasmo*GEM growth phenotype | Construct designs | Transfected constructs | Recovered transfectants | Positive selection (post-thaw) | Negative selection |
| --- | --- | --- | --- | --- | --- |
| Essential | 31 | 14 (45.2%) | 6 (42.9%) | 2 (33.3%) | 2 (100%) |
| Slow | 392 | 173 (44.1%) | 108 (62.4%) | 74 (68.5%) | 51 (68.9%) |
| Dispensable | 34 | 15 (44.1%) | 12 (80.0%) | 8 (66.7%) | 5 (62.5%) |
| No orthologue | 65 | 32 (49.2%) | 26 (81.3%) | 9 (34.6%) | 8 (88.9%) |
| **Total** | **522** | **234 (44.8%)** | **152 (65.0%)** | **93 (61.2%)** | **66 (71.0%)** |

Several additional lines were generated separate from the automated transfection pipeline. Constructs targeting PfRH3 (PF3D7_1252400) and PfEBA165 (PF3D7_0424300) were included as controls as both have previously been reported as pseudogenes in *P. falciparum*, and previous disruptions of these genes resulted in non-significant growth phenotypes during the blood stages of the parasite life cycle (Taylor, Triglia et al. 2001, Triglia, Thompson and Cowman 2001, Proto, Siegel et al. 2019). Another locus, PfPARE (PF3D7_0709700) was also included as a control since its disruption is also thought to be dispensable in blood stages based on the growth phenotypes of parasite lines with a missense mutation in the protein catalytic site (Istvan, Mallari et al. 2017), although minor growth defects resulting from disruption of the PfPARE locus and tagged with reporter genes have also been reported (Hoshizaki, Jagoe and Lee 2022). Multiple deletion lines with different barcode modules were generated for both PfRH3 and PfPARE to provide multiple controls as well as approaches to account for residual variation and experimental robustness. While the majority of deletions were made in the NF54 background, an additional three deletion lines targeting the PfPARE locus were also made in Dd2, 3D7 and GB4 strains as controls for the effect of parasite genetic background.

### Assessment of parasite growth using individual lines in arrayed screens

The effect of deleting individual genes on parasite growth was measured using both an arrayed approach, where deletion lines were cultured individually and growth monitored using flow cytometry every 48h, and also using mixed pools containing all deletion lines, where relative growth could be quantitated using BarSeq (Bushell, Gomes et al. 2017). In both assays parasite cultures were sampled every 48h, grown in the same incubators and cultured for five cycles (for more details, see Methods).

To select the most relevant line to be used as a control, growth rates for the control lines PfRH3 (variant 1), PfRH3 (variant 2), PfPARE (NF54) and PfEBA165 were tested initially along with the NF54 wild type line. All these control lines had growth rates that overlapped with the wild-type NF54 line, although with some differences in the precise mean Relative Growth Rate (RGR) growth estimates (**Supplementary Figure 2)**. Similarly, wild-type Dd2 and PARE [Dd2], and wild-type 3D7 and PARE [3D7], also had non-significant growth differences, although both wild-type Dd2 and PARE [Dd2] grew faster than 3D7 and NF54 lines, consistent with previous observations (Reilly, Wang et al. 2007, Murray, Stewart et al. 2017). The lack of significant difference between NF54 PfRH3, PfPARE and PfEBA165 knockouts relative to NF54 meant that in theory any could be used as controls to compare relative growth rates with other deletion lines, however PfRH3 was selected because it had the closest average RGR to WT NF54.

Deletion lines were scored as either slow (if the deletion line had a relative growth rate [RGR] significantly lower than the PfRH3 control lines with no overlapping confidence intervals), non-significant (if the deletion line and the PfRH3 control lines had overlapping confidence intervals), or fast (if the deletion line had a RGR significantly higher than the PfRH3 control lines and with no overlapping confidence intervals). Estimates of growth rates from wells that had been initially seeded with synchronised cultures are summarised in **Fig 1a** and show that 51/65 (78.4%) lines had slow growth phenotypes, 11/65 (16.9%) lines had non-significant growth phenotypes, whereas two lines, PfPARE [Dd2] and PfACS9 (PF3D7_0215000) had fast growth phenotypes with EBA175 [Dd2] (PF3D7_0731500) closely bordering significance for fast growth. By contrast, when the same arrayed assays were conducted using non-synchronised starting cultures, only 24/70 (34.3%) lines had a slow growth phenotype, while 41/70 (58.6%) were non-significant and 5/70 (7.1%) had a fast growth phenotype (shown in **Fig 1b**). Both PfACS9 and PARE [Dd2] had high growth phenotypes in both synchronous and asynchronous conditions. All the control lines in synchronous culture and asynchronous culture had peak distributions within expected ranges and concordant with genetic background (**Fig 1c – 1d**), suggesting robustness in these experiments. There was a significant positive correlation between RGR estimates in synchronous and asynchronous culture (*r* = 0.60, *P* < 0.001), shown in **Fig 1e**, however some lines such as EBA175 [Dd2] had negative residuals that were greater than two standard deviations from the PfRH3 control lines.

**Figure 1.**
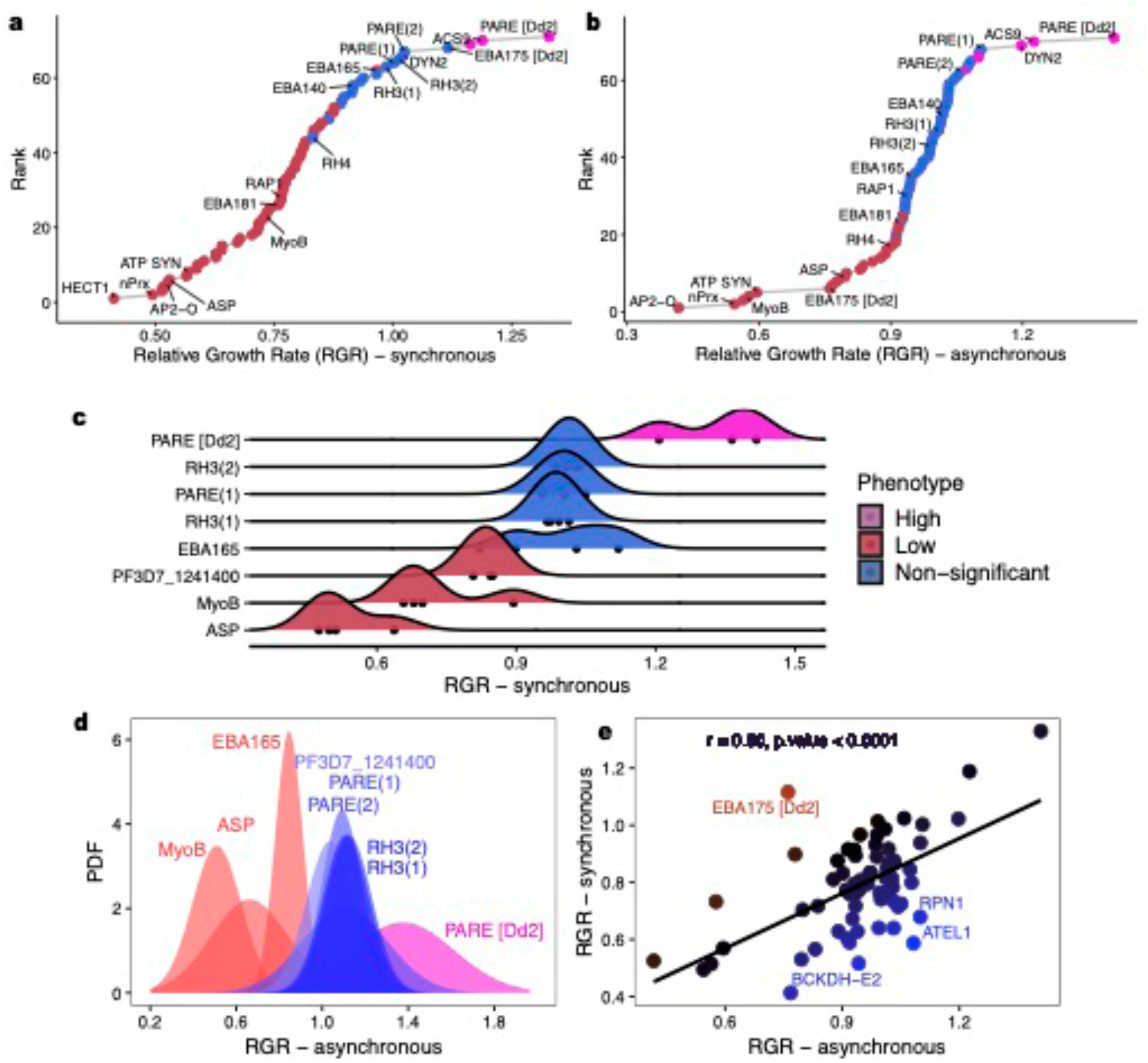
Growth phenotypes of *Plasmodium falciparum* deletion lines in static culture using arrayed approaches. *P. falciparum* lines cultured in static culture starting with either synchronous culture (a) or asynchronous culture (b). Parasite proliferation was assessed using flow cytometry. Significance of the individual gene deletions on parasite growth were then estimated by calculating 95% confidence intervals for the mean relative growth rates. PfRH3 (variant 1) and PfRH3 (variant 2) were used to determine the boundaries of significance. Several controls were used for both the synchronous approach (c) and asynchronous method (d). There was a significant positive correlation between both synchronous and asynchronous approaches (e).

### Assessment of parasite growth using pooled lines by BarSeq

For the pooled approach, the deletion lines were grown in a single mixed pool containing all 65 lines mixed in equal proportions. This pool was created once, split into triplicates, then sampled over sequential growth cycles. Growth was assessed by measuring mean fold changes in the proportion of each barcode every 48h over five cycles. Growth rates for each deletion line were significantly correlated across the three replicates using multiple linear regression (Adjusted R-squared = 0.60, *P* < 0.001), which suggested that there were minimal batch effects (shown in **Supplementary Figure 3)**. Two approaches were used to analyse the growth data of the mixed pools, differing primarily in how the confidence intervals were calculated. The first method BarSeq (RGR) was based on estimations of the per 48h cycle fold changes in growth rates for each deletion line normalized to the average per 48h cycle fold change of PfRH3 (variant1) and PfRH3 (variant2) controls (shown in **Fig 2a**); this approach is most comparable to RGR for the arrayed mutants. In the second approach BarSeq (MFC), mean fold changes in growth rates were calculated using a regression of the cumulative fold changes independent of the two PfRH3 control variants but with the expectation that parasite lines with similar growth profiles would overlap. Growth phenotypes were scored as slow, non-significant and fast following a similar approach described using the arrayed screen (summarized in **Fig 2b**). The results using the BarSeq (MFC) approach show that 46/65 lines (70.8%) had slow growth phenotypes, 19/65 (29.2%) had non-significant growth phenotypes, whereas there were no fast growth phenotypes scored. More non-significant phenotypes were observed with the BarSeq (RGR) method, where 34/65 lines (52.3%) had slow growth phenotypes, 31/65 (47.7%) had non-significant growth and similarly no fast growth phenotypes were scored. Controls PfEBA165 and PfPARE both had non-significant growth rates relative to PfRH3 (variant 1) and PfRH3 (variant 2), consistent with the arrayed approach. A summary of MFC scores plotted in ranked order is shown in **(Supplementary Figure 4)**. Growth rates generated using BarSeq (MFC) and BarSeq (RGR) were positively correlated (*r* = 0.93, *P* < 0.0001) as shown in **(Supplementary Figure 5)**. There was a significant positive correlation between the arrayed and pooled screen (*r* = 0.54, *P* < 0.0001) as shown in **Fig 2c**, with all the differences in phenotype calls at the boundaries of a gradient of either low/non-significant or non-significant/high (shown in **Fig 2d**). This also suggests that there were no significant interactions in proliferation of deletion lines in mixed pools under these growth conditions, which might potentially confound growth rates as shown for wild strains of Dd2 and HB3 (Wacker, Turnbull et al. 2012). Growth measurements using the pooled approach were repeated with a smaller subset of genes over multiples rounds, and these were consistent with previous observations. BarSeq therefore appears to be a robust method for measuring growth of *P. falciparum* lines in a scalable manner and has considerable benefits of throughput over an arrayed approach.

**Figure 2.**
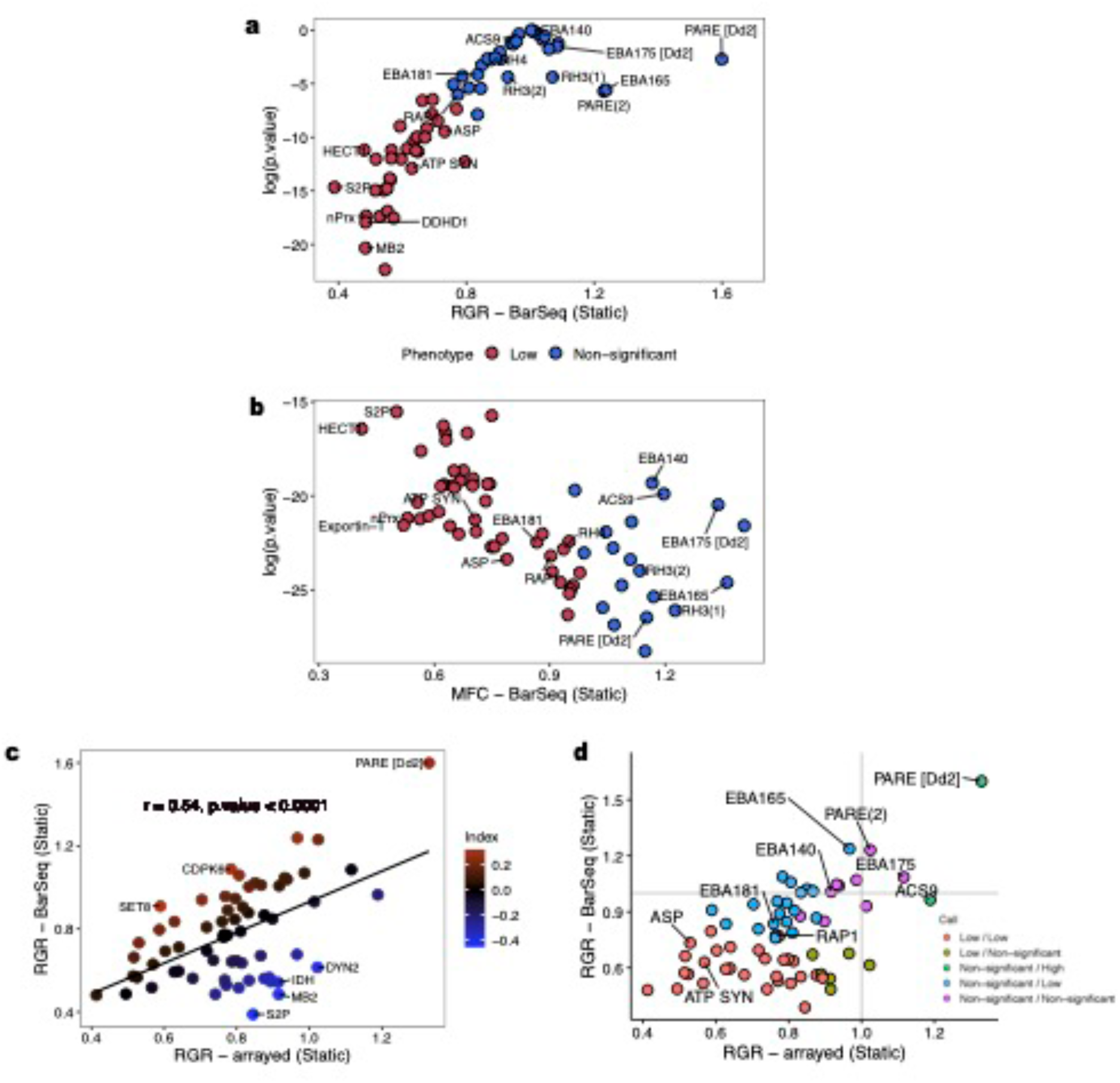
Growth phenotypes of *Plasmodium falciparum* deletion lines in static culture using pooled approaches and BarSeq. *P. falciparum* lines were synchronized, pooled and sampled for growth in static conditions. Parasite proliferation was assessed using barcode quantitation and next generation sequencing. Significance of the individual gene deletions on parasite growth were then estimated by calculating 95% confidence intervals for either the mean relative growth rate (a) or mean fold change estimate (b). PfRH3 (variant 1) and PfRH3 (variant 2) were used to determine the boundaries of significance. There was a significant positive correlation between arrayed and pooled growth approaches (c-d).

### Validation of growth phenotypes

Growth rates and phenotypes from both the arrayed and mixed pool approach were first validated by comparing them with previously published data from the large-scale *Plasmo*GEM and *Piggy*Bac screens in *P. berghei* and *P. falciparum* respectively (Bushell, Gomes et al. 2017, Zhang, Wang et al. 2018). Growth rates measured using both the arrayed and mixed pool approaches were significantly positively correlated with the *Piggy*Bac mutagenesis fitness score (MFS) and *Plasmo*GEM relative growth rates (shown in **Table 2**). Correlations in RGR were more similar to *Piggy*Bac mutants (which were made in *P. falciparum*), than *Plasmo*GEM mutants (which were made in *P. berghei*). The proportion of mutants with a particular qualitative growth phenotype were also similar across studies (summarised in **Supplementary Figure 6**). This therefore suggested that there was broad agreement between the studies, although discordant matches also existed, most of which bordered on the margins of whether a line was scored as slow or non-significant vs non-significant or fast. There were no obvious GO terms enriched for either the discordant or concordant genes. These differences might underlie biological differences between species, or technical differences in the environment or methodology of the assays.

**Table 2.** Comparison of methods used to measure growth of *Plasmodium* mutants. Parasite gene deletions that were orthologous between all the methods (total of 53 orthologues in *P. berghei*) were initially determined and a correlation test done to assess significance in association.

| Method | BarSeq (RGR) | BarSeq (MFC) | PiggyBac<br>(MFS) | PlasmoGEM | PlasmoGEM<br>(RGR > 0.5) |
| --- | --- | --- | --- | --- | --- |
| Arrayed (RGR) | 0.64 ($P < 0.001$) | 0.57 ($P < 0.001$) | 0.25 ($P = 0.02$) | 0.08 ($P = 0.58$) | 0.39 ($P = 0.01$) |
| BarSeq (RGR) | 1 | 0.93 ($P < 0.001$) | 0.41 ($P = 0.001$) | 0.27 ($P = 0.05$) | 0.40 ($P = 0.01$) |
| BarSeq (MFC) | 0.93 ($P < 0.001$) | 1 | 0.41 ($P = 0.001$) | 0.30 ($P = 0.03$) | 0.41 ($P = 0.01$) |
| PiggyBac (MFS) | 0.41 ($P = 0.001$) | 0.41 ($P = 0.001$) | 1 | 0.43 ($P = 0.001$) | 0.30 ($P = 0.06$) |
| PlasmoGEM | 0.27 ($P = 0.05$) | 0.30 ($P = 0.03$) | 0.43 ($P = 0.001$) | 1 | - |

As an additional validation, completely new constructs were generated targeting three genes (PfHECT1/PF3D7_0628100, PfnPrx/PF3D7_1027300, PF3D7_0318500) which resulted in a slow growth phenotype in the initial RGR assessment. Growth estimates of these repeat deletion lines were then determined using the arrayed approach (as described in Methods). Control deletion lines PfRH3 (variant 1) and PfRH3 (variant 2) were included, as well as lines with deletions in PfEBA181 and PfAP2-O which had a slow growth phenotype as shown in this study. Growth phenotypes of these repeat deletion parasite lines (PfHECT1, PfnPrx, and PF3D7_0318500) were all significantly lower than the two PfRH3 controls, consistent with previous observations (shown in **Supplementary Figure 7**).

### Comparing parasite growth between static and shaking conditions

*Plasmodium* blood-stage parasites encounter diverse environmental conditions *in vivo* such as changes in blood pressure and volume, variation in circulating metabolites, variations in host RBC types and ages, as well as different tissue such as the spleen, liver, bone marrow, all of which might have an effect on parasite growth (Kho, Qotrunnada et al. 2021). Growth is therefore not necessarily a fixed phenotype for a given strain. The advantage of BarSeq is that it provides a scalable method that can be used to assess growth of multiple lines across many different conditions.

To explore the utility of BarSeq for such studies, we first tested the effect of mechanical stimuli on parasite growth by maintaining mixed pools in shaking and static conditions in parallel, and generating RGR and MFC estimates from three biological replicates as described above. There was a significant correlation between the biological replicates (Adjusted R-squared = 0.80, *P* < 0.001) with less variation in shaking conditions compared to static culture (shown in **Supplementary Figure 8**). In shaking culture, a total of 44/65 (67.7%) lines had slow growth phenotype whereas 21/65 (32.3%) lines were non-significant based on RGR estimates. No fast growers were scored using RGR in shaking conditions (shown in **Fig 3a**). A higher proportion of slow growth phenotypes was observed with the MFC method 58/65 (89.2%) whereas 7/65 (10.8%) lines were non-significant (shown in **Fig 3b**). There was a significant correlation between RGR and MFC estimates from shaking culture (*r* = 0.927 *P* < 0.0001) as shown in **Supplementary Figure 9**.

**Figure 3.**
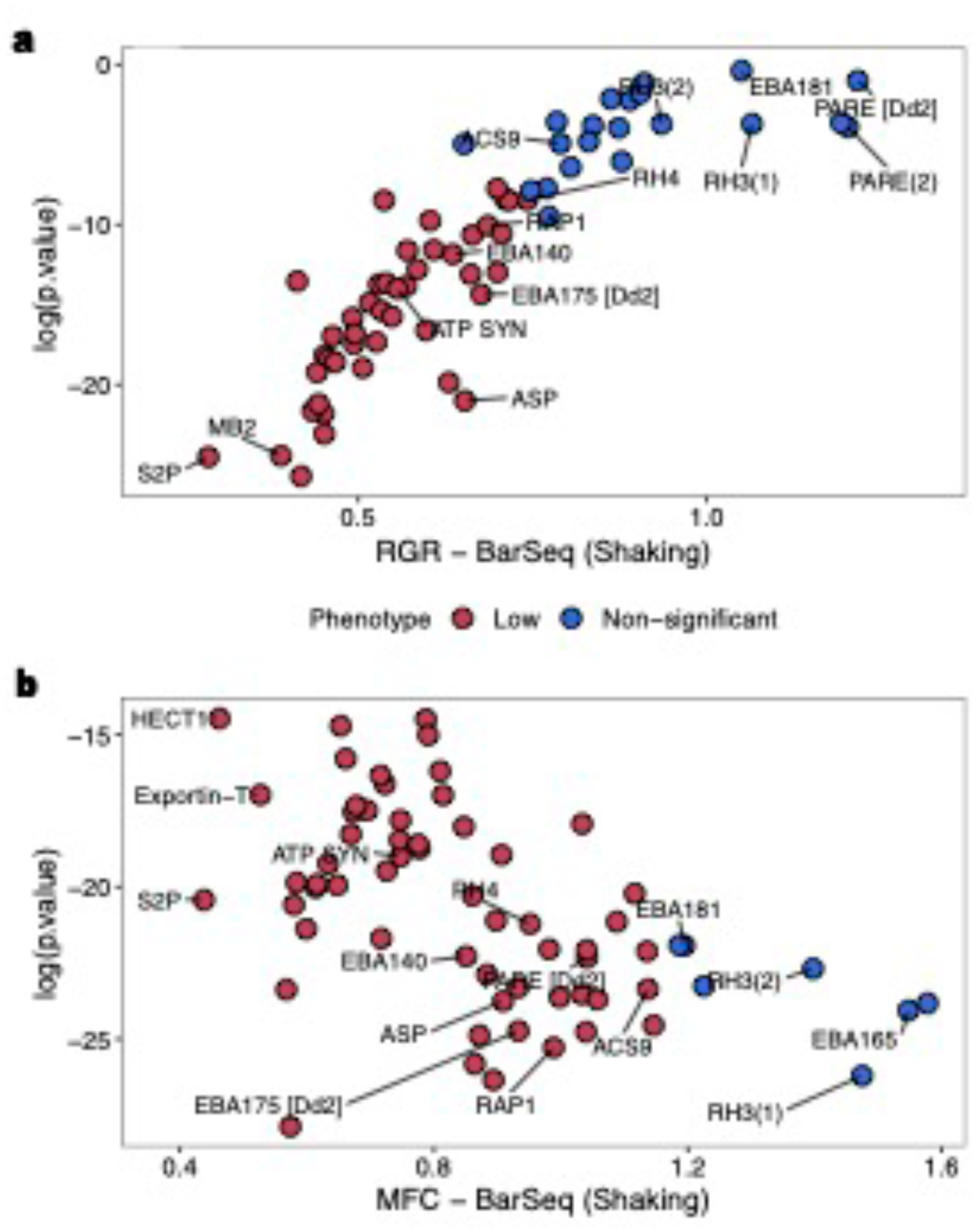
Effect of shaking on parasite growth. *P. falciparum* lines were synchronized, pooled and sampled for growth in shaking conditions. Parasite proliferation was assessed using barcode quantitation and next generation sequencing. Significance of the individual gene deletions on parasite growth were then estimated by calculating 95% confidence intervals for either a mean relative growth rate (a) or mean fold change estimate (b).

Formal comparisons of mean fold changes in growth rates in either shaking or static culture (shown in Figure **4a**) were performed using cumulative fold changes for each deletion line, fitted with a regression line using three independent replicates per cycle. This regression was then used to generate an index we have referred to as the shaking vs static index (SSIGR); deletion lines that grow better in static conditions will have a negative SSIGR index whereas deletion lines that grow better in shaking conditions will have a positive SSIGR index, and this can be exponentiated. PfRH3 (variant 1) and PfRH3 (variant 2) were used as controls to estimate 95% confidence interval boundaries for assignment of phenotypes, and again the other controls, PfPARE and PfEBA165, fell within these boundaries **(Fig 4b**). A total of 1/65 (1.5%) deletion lines had a high SSIGR, 20/65 (30.8%) had non-significant SSIGR and 44/65 (67.7%) had low SSIGR estimates, meaning they grew faster in static conditions. The most extreme lines were all deletions in the Erythrocyte Binding Ligands (EBLs), notably PfEBA181 with high SSIGR and both PfEBA140 and EBA175 [Dd2] with low SSIGR, suggesting that shaking significantly affects growth rates in these invasion-related deletion lines and hence potentially modulates gene function of these ligands.

**Figure 4.**
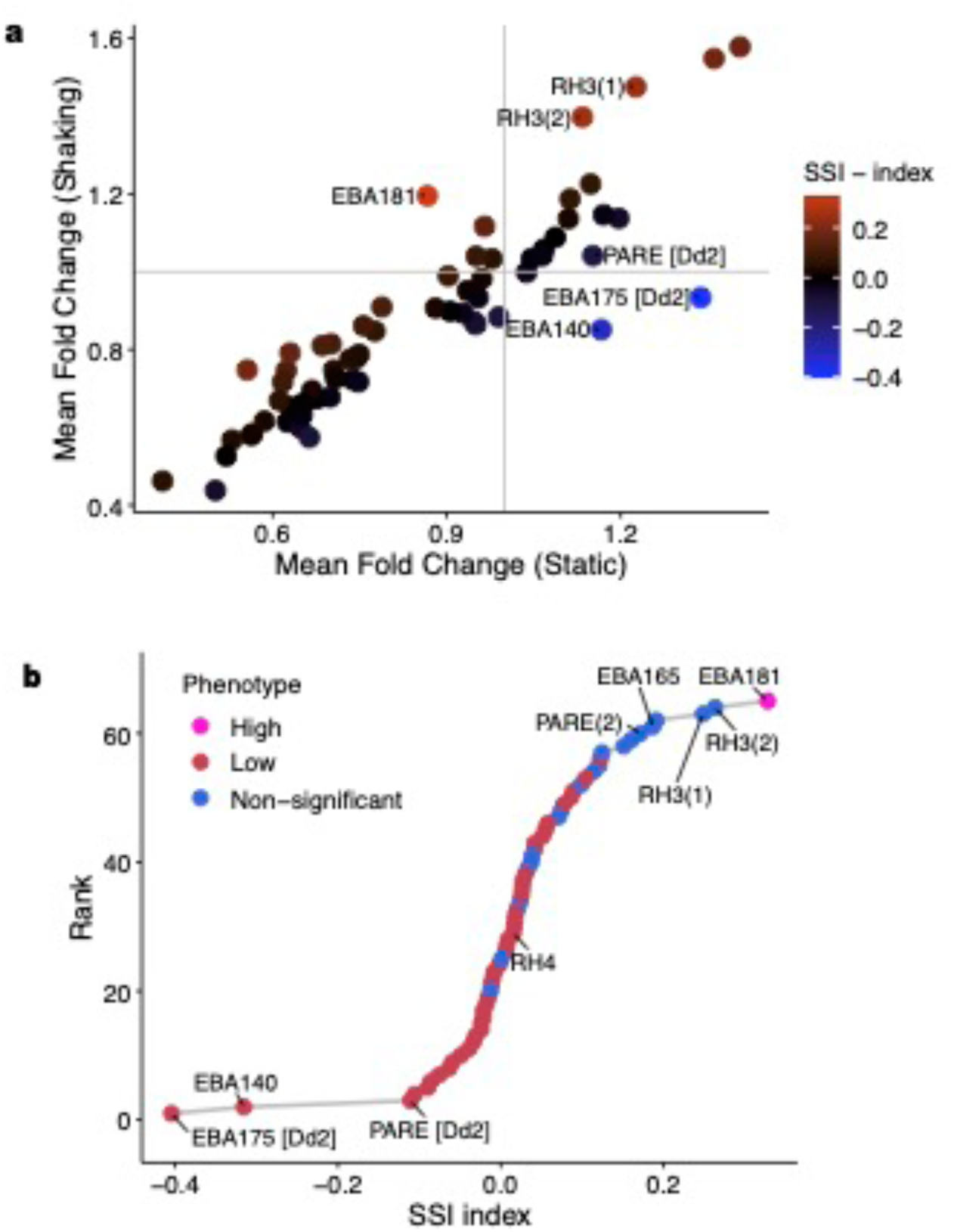
Comparison of shaking and static culture on parasite growth. Growth of *P. falciparum* deletion lines in shaking and static cultures were also compared in parallel (a). Differences in parasite growth were estimated using the shaking vs static index (SSI_GR_). Control lines PfRH3 (variant 1) and PfRH3 (variant 2) were used to determine boundaries for significance. Phenotypes were scored as either low, non-significant or high (b).

### Using arrayed and BarSeq approaches to quantitate DNA replication

To further explore the potential for using BarSeq in more complex phenotyping screens, we applied it to estimate DNA replication capacity of each strain. Variations in merozoite number within a single schizont exist between *Plasmodium* species, strains and even within clonal populations (Simon, Stürmer et al. 2021). Merozoite number can be estimated by using flow cytometry to quantitate DNA content in late stage schizonts that have been arrested just before egress using the Protein Kinase G inhibitor, Compound 2 [(Donald, Zhong et al. 2006); see Methods]. As with growth assays, we assessed DNA content in the deletion lines using both an arrayed and mixed pool/BarSeq approach, with the arrayed approach being carried out in four batches.

Based on the arrayed approach (shown in **Fig 5a**), C2-arrested schizonts from a total of 8/70 (11.4%) lines had significantly higher DNA content than the PfRH3 controls, 18/70 (25.7) had lower levels, and 44/70 (62.9%) were non-significantly different. The deletion lines with the lowest DNA content were EBA175 [Dd2], PfACS9 and PARE [Dd2]. It has been previously reported that wild type Dd2 makes an average of 18 daughter merozoites per schizont (Reilly, Wang et al. 2007); this was consistent with our observations of both PARE [Dd2] and wild type Dd2 DNA content, both of which were consistently significantly lower than wild type NF54. High levels of kurtosis, a parameter which describes the distribution of the data around the mean estimate, were also observed for PARE [Dd2] and EBA175 [Dd2]. Deletion of BCKDH-E2 resulted in a much broader spread of DNA content compared to other strains, which could also indicate a dysregulation of replication. Several controls were included, and these overlapped with the non-significant distribution of the RH3 controls. The distribution of the DNA content for the individual deletion lines is summarised in **Supplementary Figure 10**.

**Figure 5.**
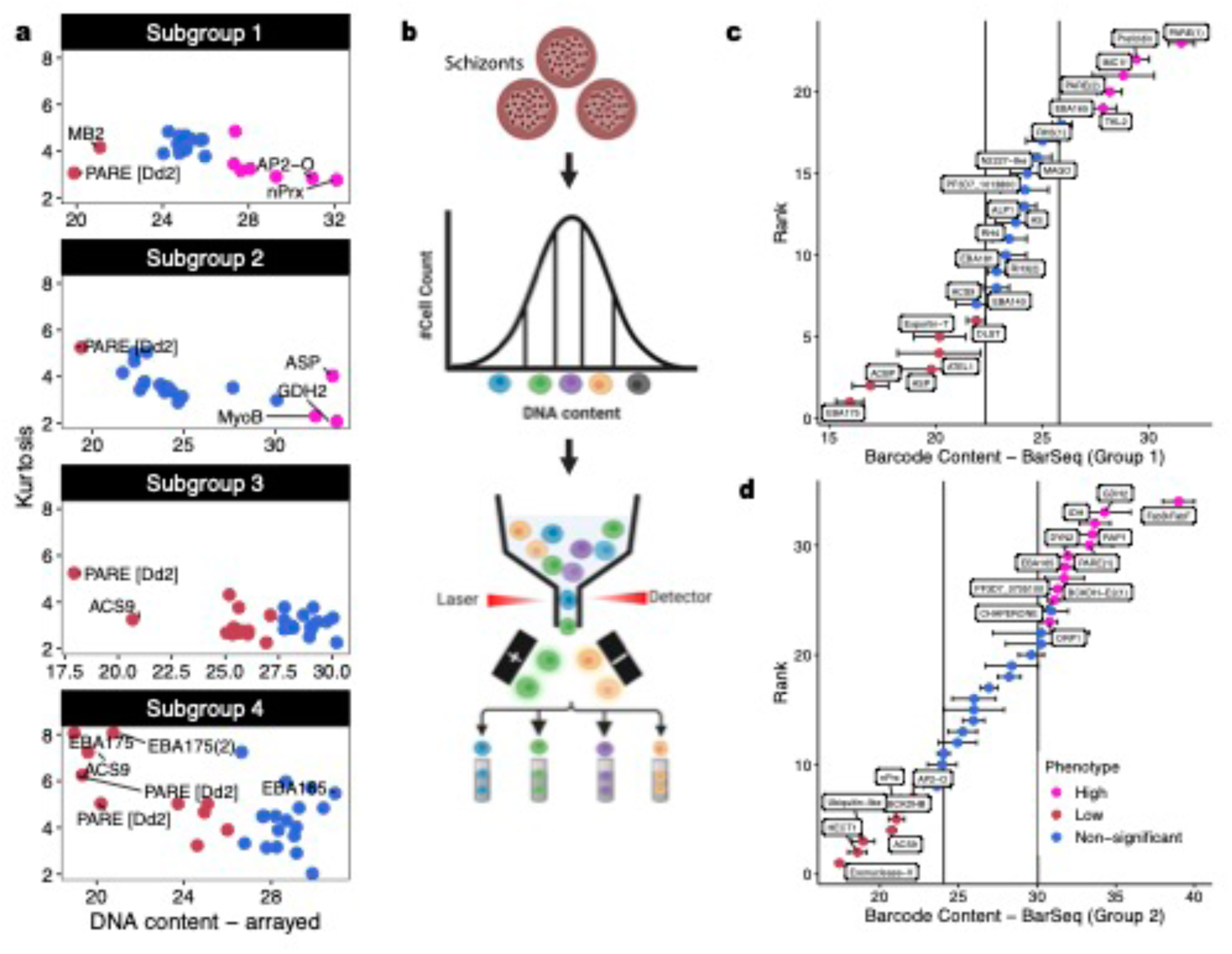
Estimates of DNA content within mature schizonts of *Plasmodium falciparum* deletion lines using arrayed, flow sorting and BarSeq approaches. Parasite DNA and barcode content was quantitated using either an arrayed or mixed pool BarSeq approach. For the arrayed approach, mature schizonts were stained with Hoechst 33342 and quantitated using flow cytometry (a). The PfRH3 lines were used as controls to determine boundaries of significance with phenotypes scores either as low, non-significant or high. For the mixed pool method, mature stage parasites were acquired on a cell sorter and their distribution subdivided into quintiles (b). Schizont populations from each window were then sorted and quantitated using next generation sequencing. A frequency distribution was used to calculate the average barcode content for each deletion line (c-d). The PfRH3 lines were used as controls to determine boundaries of significance with phenotypes scores either as low, non-significant or high.

BarSeq experiments were performed using two mixed pools of deletion lines, each containing the same six controls – PfPARE, PfACS9, PfRH3 variant 1, PfRH3 variant 2, PfExportin-T (PF3D7_1315900), PfEBA165, along with 17-28 other randomly selected deletion lines. The distribution of the relative DNA fluorescence units of the arrested schizont mixed parasite pools followed a normal distribution (**Supplementary Figure 11**). The mixed pool distribution was then subdivided into quintiles, which were used as gates for flow sorting (shown in **Fig 5b**). DNA was then extracted from each sorted population, and the number of barcodes for each line within each quintile quantitated using BarSeq. Reads were standardized using relative fluorescence units to determine the number of infected red cells per deletion line within each quintile (shown in **Supplementary Figure 12**). The barcode distribution of each deletion lines was scored as either low (if the deletion line had barcode content significantly lower than the PfRH3 control lines with no overlapping confidence intervals), non-significant, or high (if the deletion line had barcode content significantly higher than the PfRH3 control lines and with no overlapping confidence intervals). In group 1, a total of 6/23 (26%) deletion lines had significantly lower barcode content than the PfRH3 controls, 5/23 (21.7%) had higher barcode content and 12/23 (52.2%) were non-significant (**Fig 5c**). In group 2, there were 11/34 (32.3%) low, 7/34 (20.6%) high and 16/34 (47.1%) non-significant deletion lines (**Fig 5d**).

A total of 53 lines were assayed using both the arrayed and BarSeq approach; Overall, there was ∼40% concordance between the phenotypes of the two approaches (shown in **Supplementary Figure 13**), however the correlation between these two methods was higher for mutants with estimated DNA content < 24 (*r* = 0.91 *P* < 0.0001) as shown in **Supplementary Figure 14**. Several deletion lines such as PARE [Dd2], EBA175 [Dd2] and PfACS9 had concordant phenotypes in both arrayed and pooled methods, suggesting a genetic basis for the regulation of DNA content. There was also a consistent negative correlation between DNA content and growth (shown in **Supplementary Figure 15**), in keeping with the known phenotypes of the Dd2 control strain, which grows faster than NF54, but produces fewer merozoites.

### Using BarSeq to track progression of mutants through the intraerythrocytic developmental cell cycle (IDC)

Once inside RBCs, *P. falciparum* parasites undergo an intricate programme of growth and replication through ring, trophozoite and schizont stages, leading to the formation of daughter merozoites which then burst out and infect new RBCs. This intraerythrocytic developmental cycle (IDC) is tightly regulated and its length varies both within and between *Plasmodium* species (Reilly, Wang et al. 2007, Reilly Ayala, Wacker et al. 2010), but the genes that control it are only imperfectly understood (Paul, Miliu et al. 2020). To apply BarSeq to this problem, we used flow sorting based on RNA and DNA fluorescence to separate parasite pools into specific developmental stages – RNA transcription peaks in trophozoite stages, whereas DNA replication only occurs in late trophozoite and schizont stages (shown in **Fig 6a**). Pools of parasites were tightly synchronised, sampled at multiple points over a 48h time course, and at each time point, the parasite population was sorted into the same gates. The relative distribution of each deletion line between gates was then quantitated using BarSeq.

**Figure 6.**
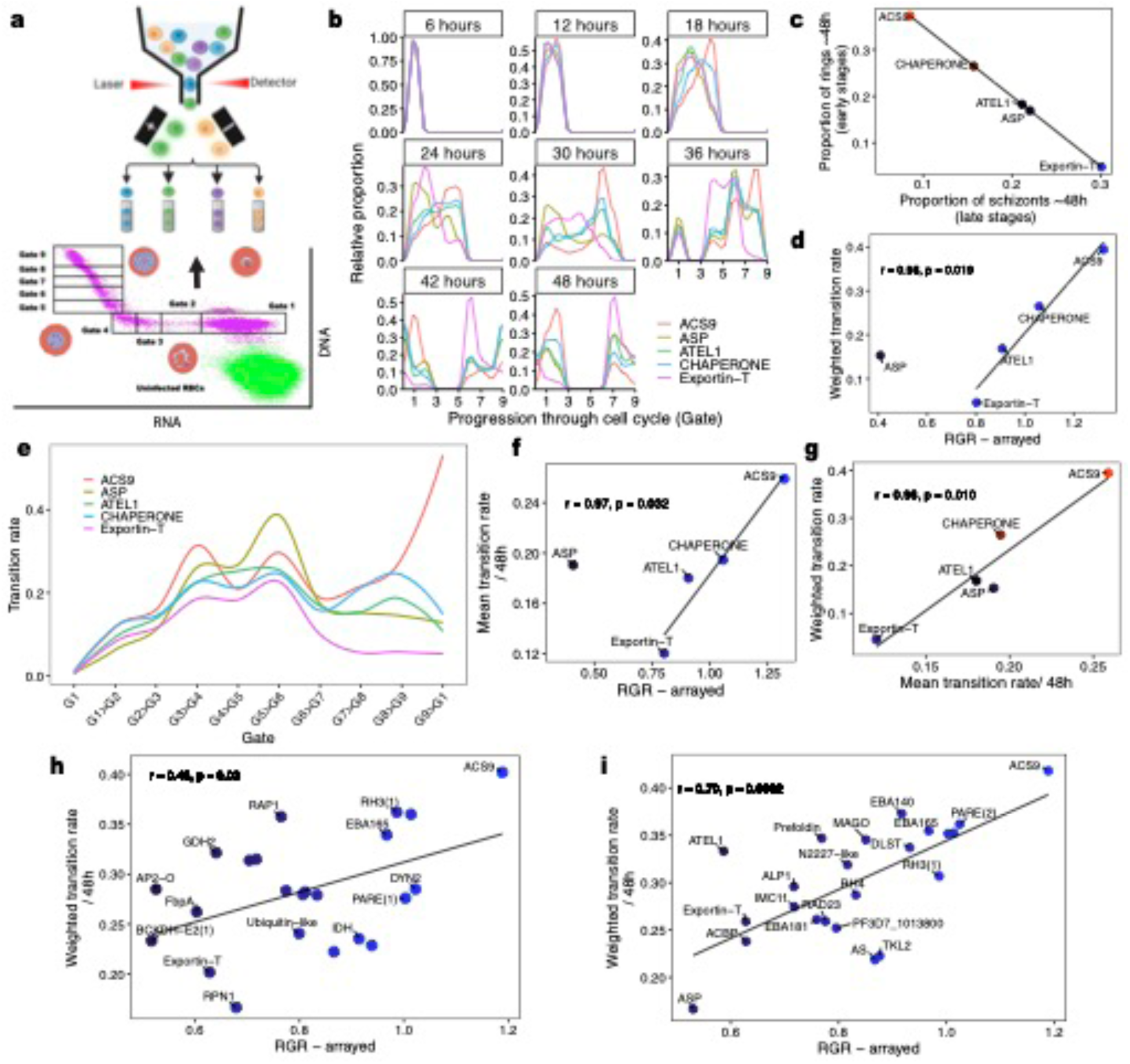
Intraerythrocytic developmental cell cycle dynamics of *Plasmodium falciparum* deletion lines determined using flow sorting and BarSeq. Synchronous parasite cultures were pooled and sorted based on their RNA/DNA content using a BD influx cell sorter (a). The relative proportion of reads of each deletion line was then assessed over a 48h time course (b). The rate of transition was then determined using a weighted approach (c-d). A second approach using a compartmental model was also used to determine the transition rate per hour between each of the gates (e). A mean transition across the cycle was also determined from this model and is positively correlated with growth (f). Both the weighted approach as well as the modelling approach had a positive correlation (g). Similar weighted transition rates were also observed for the larger pools, and these were all significantly correlated with growth and associated with genetic background (h-i).

Three independent runs were performed as described in more detail in the Methods section. The first experiment had a pool of five deletion lines (PfACS9, PfExportin-T, PfChaperone (PF3D7_0920100), PfASP and PfATEL1 (PF3D7_0212900) and was set up as the proof of principle. The changes in barcode proportions between multiple gates sampled every 6h over a 48h time course are summarized in **Fig 6b**, and from these trends can be inferred. At 6h, all deletion lines are within the earliest gate, however a bimodal distribution emerges at 12h with PfACS9 and PfChaperone being more advanced in the cell cycle. By 18h, a trimodal distribution is present with PfACS9 the most advanced strain, PfChaperone is intermediate, whereas PfExportin-T, PfASP and PfATEL1 are slowest. By 36h, PfACS9 is further advanced in the cell cycle whereas PfExportin-T is the slowest and by 48h, parasites have started re-invading new RBCs with PfACS9 further advanced in the cell cycle followed by, PfChaperone, PfATEL1/ PfASP and PfExportin-T in that order.

We explored two methods of quantitating the rate of transition of these deletion lines though the IDC. The first approach following a ‘before and after’ approach is primarily designed to identify differences in progression after the full time-course. At 48h, PfACS9 had the highest proportion of reads in the early gates, indicating completion of one cycle and beginning of another, whereas PfExportin-T had a higher frequency of reads in the late stages (shown in **Fig 6c**). A weighted mean of the barcode counts was then calculated using ring stage parasites and showed that PfACS9 was the quickest to advance whereas PfExportin-T was the slowest (shown in **Fig 6d**). There was also a significant positive correlation between the weighted transition rate and growth (as measured using the arrayed approach/RGR) (*r* = 0.98 *P* < 0.019), although PfASP was an outlier. The second approach used a compartmental model to simulate the transition of reads between gates per hour (shown in **Supplementary Figure 16**). Each compartment records the proportion of sequencing reads observed at time *t* in that gate on the assumption that the rates of transitions between the gates are constant for a particular deletion line. Results are shown in **Fig 6e**. As seen using the ‘before and after’ approach, PfACS9 transitions fastest between gate 3-4, and gate 5-6 and with a fast transition in the late stages. PfASP transitions slowest in the earliest stage (gate 1-3), whereas PfExportin-T generally transitions slowest. All the lines appeared to transition slowest in gates 4-5, and gates 6-7. A mean transition rate from the modelled estimates was also calculated (shown in **Fig 6f**) and again shows a positive correlation with arrayed growth (RGR) (*r* = 0.97 *P* < 0.032), with ASP again an outlier. Both the weighted transition rate and the mean transition modelled rate were positively correlated (*r* = 0.96 *P* < 0.010) as shown in **Fig 6g**.

The weighted approach was then used to determine the transition rates in two larger pools of deletion lines (shown in **Fig 6h-i**). There was a positive correlation between the weighted transition rate and arrayed growth, as well all RGR from BarSeq data (shown in **Supplementary Figure 17**-18), and multiple deletion lines were consistent across all the IDC experiments such as PfACS9 which transitions fastest through the IDC in all pools of deletion lines tested.

## Discussion

We generated a medium-scale library of targeted *P. falciparum* knockout lines using a custom CRISPR/Cas9. This is only the fourth targeted genetic study of this scale in the major human malaria pathogen, comparable to previous studies of kinases (Solyakov, Halbert et al. 2011), exported proteins (Maier, Rug et al. 2008) and unknown proteins on chromosome 3 (Kimmel, Schmitt et al. 2023). Rather than targeting a specific group of genes, or genes on a specific chromosome, as in previous studies, the gene list was selected to enrich for slow growth phenotypes, based on a previous targeted screen in the rodent model *P. berghei* (Bushell, Gomes et al. 2017, Zhang, Wang et al. 2018). Unlike previous studies however, we focussed on scaling up phenotyping by using BarSeq to assess multiple parasites growing simultaneously in mixed pools. BarSeq was applied to quantitate growth rates *in vitro,* analogous to the way it was applied *in vivo* in *P. berghei,* but also in two completely novel ways – to quantitate DNA multiplication, and to follow cell cycle transitions.

There was overall a positive correlation between parasite growth rates in our study and those in the *Plasmo*GEM and *Piggy*Bac screens, even though the screens were carried out using very different methodologies. 73.6% of the deletion lines in this study using BarSeq had concordant growth phenotypes with orthologues in the *Plasmo*GEM screen, despite the screens being carried out in different *Plasmodium* species. Discordant phenotypes could be the result of genuine differences in biology between species but could also be driven by methodological differences. In the *Plasmo*GEM screen the full ORF was replaced by a selectable marker, whereas in our *P. falciparum* pipeline, the marker deleted only a section of the ORF, which could result in expression of partial gene fragments that might provide partial function. We mitigated this by targeting the 5’-end of the gene to introduce premature stop codons upstream of the sequence and hence increase the probability of generating non-functional truncated genes, but it remains a possibility that in some cases our disruptions were not complete. Similarly, potential differences could exist between culture *in vivo* system as used in the *Plasmo*GEM screen and the *in vitro* system used in this study. Several other large genome-scale screens have reported a higher abundance of dispensable genes in *in vitro* culture and free-living organisms under laboratory conditions (Winzeler, Shoemaker et al. 1999, Giaever, Chu et al. 2002, Sidik, Huet et al. 2016). There was higher correlation in growth estimates with the *Piggy*Bac transposon data, which was carried out *in vitro* in *P. falciparum,* so concordance would be expected to be higher. Discrepancies here could be driven by inaccessibility of certain regions of the genome to TTAA insertion, incomplete saturation or in other cases insertions at multiple sites within the genome, leading to complex phenotypes.

Some of these differences may reveal new biology. For example, the *Plasmo*GEM screen only identified a single knockout that resulted in faster growth – deletion of the AP2G locus which inactivates gametocyte switching and therefore results in fewer gametocytes being produced, which are a dead-end stage during intraerythrocytic development (Sinha, Hughes et al. 2014, Bushell, Gomes et al. 2017). We observed consistently higher growth phenotypes of an ACS9 knockout line which has not been reported in other screens. Confirmation of an intact AP2G locus in these ACS9 deletion line was further confirmed using whole genome sequence analyses (shown in **Supplementary Figure 19**). ACS9 is part of a family of fatty acyl-CoA synthetases that catalyse the formation of a thioester bond between fatty acids and coenzyme-A (CoA) forming acyl-CoA, a precursor for phosphatidic acid which is key to phospholipid metabolism (Athenstaedt and Daum 1999, Watkins 2008). Notably. Although *P. berghei* contains a single direct homologue of ACS9, this ACS9 gene has further expanded through duplication in *P. falciparum* into a family of 10 paralogs (Bethke, Zilversmit et al. 2006). This could possibly explain why it was not identified as a fast growing mutant in the *Plasmo*GEM screen, and points to differences in phospholipid metabolism between *Plasmodium* species.

Screens in other organisms have emphasised that gene function can depend heavily on the growth environment, for example in yeast cells where ∼80% of the genome is dispensable in rich-medium but nearly all of these genes, totalling 97%, revealed a growth phenotype when the cells were exposed to different and specific environments (Hillenmeyer, Fung et al. 2008). We took advantage of the high-throughput nature of BarSeq phenotyping to quantitate the effect of mechanical stimuli on parasite growth by culturing parasites in parallel in shaking and static culture and comparing growth in these two conditions by developing an SSI index. Parasites are normally grown in static culture; however, several publications have demonstrated improved growth when parasites are cultured in shaking conditions, potentially due to increased random distribution of merozoites leading to fewer erythrocytes being invaded by multiple merozoites, and more equal distribution of nutrients and metabolites through the culture medium (Butcher 1979, Butcher 1981, Zolg, MacLeod et al. 1982, Allen and Kirk 2010). Comparing the SSI index of wild type parasite strains provided evidence for strain-specific effects, with NF54 parasite lines having a positive index and hence growing better in shaking culture whereas Dd2 had a negative index and grows better in static culture. All the control lines (RH3 variant 1, RH3 variant 2, EBA165, PARE) had a similar phenotype to wild type NF54, whereas many of the gene deletions had a slight negative SSI index meaning they had a growth advantage in static culture, suggesting that the effect of specific genotypes on parasite survival could be modulated by specific environmental factors, and that shaking culture can be a way to tease out more subtle growth phenotypes not detected under static conditions. Deletion lines at the extreme end of this distribution were all members of the EBL family, specifically EBA181, EBA140 and EBA175 [Dd2] (Wright and Rayner 2014, Cowman, Tonkin et al. 2017). All these ligands bind specific receptors on the RBC surface and are thought to be involved in RBC recognition, deformation and re-orientation of the merozoite during the process of invasion. Knockout of these genes in other studies has also been associated with a reduction in parasite invasion consistent with our results (Weiss, Gilson et al. 2015, Koch, Wright et al. 2017), but the more marked differences revealed in shaking conditions again suggest that varying growth conditions can exacerbate phenotypes, and that in the case of these genes, shaking conditions may influence gene function and better reproduce the situation *in vivo* where merozoites and RBCs interact while exposed to blood flow.

We also used BarSeq for the first time to compare DNA content, and hence multiplication, between strains. Differences in proliferation rates between strains Dd2 and HB3 have previously in part been associated with differences in the number of merozoites that each produce (Reilly, Wang et al. 2007), but this phenotype has never been explored at high scale. We measured the DNA content of arrested late-stage schizonts from wild type and deletion lines in both an arrayed layout using DNA staining and a pooled format which combined DNA staining with cell sorting and BarSeq. We observed lower DNA content estimates for both PARE [Dd2] and EBA175 [Dd2] than for wildtype NF54, but in line with estimates from (Reilly, Wang et al. 2007) who used microscopy to count the distribution of merozoites within 50 schizonts of wild type Dd2. The high throughput BarSeq approach by contrast allowed us to generate estimates based on counts of >10,000 schizonts allowing reductions in standard errors of estimates and identification of more subtle differences which could be significant in a large parasite biomass such as occurs *in vivo*. Phenotype scores of DNA content varied from either low, non-significant or high and there was overall a negative correlation between DNA content and growth rates, consistent with known growth and merozoite number phenotypes for Dd2 relative to NF54/3D7. Certain deletions resulted in low DNA content estimates for example ACS9, whereas deletions in other genes such as GDH2 resulted in higher estimates, suggesting multiple genes may regulate DNA content. We also assessed variability in DNA content using kurtosis of the distribution of the DNA content and observed that PARE [Dd2] and EBA175 [Dd2] generally had high kurtosis whereas other deletion lines such as BCKDH-E2 had broader variation in the final DNA content. All these might reflect variations in the replication dynamics and regulation associated with parasite genotype. The general consistency between replicates and between the two different methods, testing lines both individually and in pools, emphasize the final DNA content in mature schizonts is relatively stable and that under standard culture conditions is intrinsic to the parasite genetic background, as has been previously suggested (Reilly, Wang et al. 2007). While there was generally consistency between the arrayed and pooled approach, discrepancies were also apparent with certain genes such as AP2-O and nPrx which had significantly higher estimates in the arrayed approach. The arrayed approach uses Hoechst 33342 to directly measure DNA quantity, whereas the pooled approach relies on BarSeq quantitation, which measured the relative distribution of barcodes between sectors of the DNA content distribution, which could account for these differences or biological factors such as cellular events of merogony.

We also used BarSeq to assess differences in cell-cycle dynamics by using Hoechst and Thiazole Orange to measure DNA and RNA content and hence progression through the cell cycle. While this dual labelling approach to follow intra-erythrocytic progression has been used before (Grimberg, Erickson et al. 2008), in this case we combined it with cell sorting and BarSeq to quantitate parasite progression between cell cycle phases. Several deletion lines had different cycling rates, as measured by two different approaches, a simple start-to-end approach and a more complex compartmental transition model. Deletion of some genes such as the ACS9 locus consistently accelerated progression through the full-length cell-cycle whereas deletion of others such as Exportin-T, putative increased the IDC duration. The resolution in cell-cycle tracking was then increased by shortening the sampling window for a subset of deletion lines. ACS9 had a shorter G1 and S/M phase whereas Exportin-T, putative had longer G1 and S/M phases. Overall, there was a positive correlation between growth and the rate of completion of the cell-cycle. Previous studies using forward genetics approaches to map QTLs associated with IDC duration differences between Dd2 and HB3 strains identified potential candidates on multiple chromosomes associated with regulation of cell cycle length (Reilly Ayala, Wacker et al. 2010), which could further suggest that this trait is polygenic and that it is tightly regulated given the conservation of transcriptomic profiles between geographically diverse *P. falciparum* lines (Llinas, Bozdech et al. 2006). The BarSeq method developed here to track cell cycle dynamics of deletion lines provides a novel approach to pinpoint gene function in specific cellular processes and could be a tool in targeted genetic screens, design and identification of new drug molecules and combinations with novel cellular targets.

We generally observed stable proliferation rates both in wild type and mutant parasite lines, consistent with the fact that growth is a heritable quantitative trait. The underlying developmental and cellular processes and how these are linked in regulating these growth differences have however been poorly understood, although in some cases these have been positively correlated with sub-phenotypes such as invasion efficiencies, IDC length and merozoite numbers. We developed methods to quantify within-cycle processes of deletion lines using BarSeq. Using simple linear regression, we observed a negative correlation between growth rates and DNA content, and a positive correlation between growth rates and IDC duration. We then modelled the combinatorial effect of DNA content and IDC progression on growth rates using either the support vector regression (SVR) model or the Dense Neural Network (DNN) and observed a significant correlation with growth rates. This could potentially suggest that for some strains, the decrease in merozoite numbers is associated with a shorter cell cycle, which could therefore lead to faster growth. Other sub-phenotypes such as invasion efficiencies might also contribute to differences in overall growth phenotypes and are linked tightly with other sub-phenotypes.

In summary, we have developed new methods to generate and phenotype a significant set of *P. falciparum* deletion lines in a scalable manner using both arrayed and pooled assays. These methods are complementary, with concordant phenotypes from several comparisons, and therefore constitute a framework to pinpoint with greater accuracy the biological processes that specific genes are involved in. To our knowledge BarSeq has never been applied to phenotype either DNA quantity pr cell cycle transition in *Plasmodium* parasites. The ability to use the same barcoded deletion lines in multiple phenotypic approaches argues strongly for the inclusion of barcodes in all future genetic modification experiments, ideally using a common barcoding module to allow for the accumulation of pools across multiple different labs and research groups. Given the difficulty and slow pace of generating modified *P. falciparum* lines, even using CRISPR/Cas9 approaches, designing all generated lines to be as useable and flexible as possible, rather than generating them only for a single study, is clearly of the greatest benefit to the malaria research community. The ability to create mutations for 1% of the *P. falciparum* genome in a single lab in this manner also suggests that community-scale whole genome targeted studies should be considered, with appropriate funding support.

## Acknowledgements

Funding: This work was supported by Wellcome, grant numbers 220266/Z/20/Z (JR) and 206194/Z/17/Z (Sanger). For the purpose of open access, the authors have applied a CC BY public copyright license to any Author Accepted Manuscript version arising from this submission. The funders had no role in study design, data collection and analysis, decision to publish or preparation of the manuscript.

We thank Emily Mallet, Emma Goffin, David Adam Jackson (Sanger CGaP), Christopher Hall (Sanger core), and Sean Wright and Tristram Bellerby (Sanger DNA Pipelines) for technical support, and Theresa Feltwell, Nadia Cross and the CIMR core support teams for their invaluable logistical support.

## Author contribution

Conceptualization, A.M., O.B. and J.C.R.; methodology, A.M, M.G., T.S., W.W.; data and resource generation, A.M., M.G., M.I., R.A., S.H., R.S., G.G., F.S; validation, A.M.; formal analysis, A.M., and J.C.R.; writing – original draft, A.M., and J.C.R.; writing – review & editing, A.M., W.W., O.B., and J.C.R.; visualization, A.M., and T.S.; supervision, E.B., C.B., O.B. and J.C.R.; funding acquisition, O.B. and J.C.R. All authors read and approved the manuscript.

## Methods

### Creation of deletion plasmid constructs

An in-house automated bioinformatics pipeline was developed to generate designs of deletion constructs at scale. Two starter plasmids, pDC2-cam-coCas9-U6-yDHOODH and pCC1-hDHFR-yFCU were used as a base for the gRNA and homology repair templates respectively (Maier, Rug et al. 2008).

Prior to cloning the target constructs, frozen stocks of both pDC2-cam-coCas9-U6-yDHOODH and pCC1-hDHFR-yFCU wild-type plasmids in XL-10 competent cells (Agilent) were initially expanded by inoculation in 200mL LB broth supplemented with 100ug/mL Ampicillin and grown for 18h at 37°C overnight under shaking conditions at 200rpm. This bacterial culture was then pelleted by centrifugation at 4000rpm for 10min. Plasmid DNA was purified using a midi prep kit (Macherey-Nagel) as per the manufacturer’s instructions.

To generate donor template constructs, primers were used to amplify parasite DNA fragments (approx. 1200bp) flanking the cleavage target site at both the 5’ and 3’ ends. Short DNA adapters complementary to digested plasmids were also attached to these primers during the design process. This generates two fragments referred to hereafter as H5 and H3 homology regions. The amplicons were run through agarose gels to ensure that the right product sizes were obtained, and PCR amplicons were purified using the Min-elute 96 UF PCR purification kit (Qiagen) following the manufacturer’s instructions.

pCC1-hDHFR-yFCU base plasmids were digested using SacII and EcoRI restriction enzymes in preparation for Gibson Assembly (Gibson, Benders et al. 2008). Confirmation of successful plasmid digestion was done using agarose gel electrophoresis. The digested reaction mix was then purified prior to Gibson Assembly using a Zymo DNA clean and concentrator kit (Zymo Research) following the manufacturer’s instructions. Purified samples were quantitated using a nanodrop to determine DNA concentrations. A total of 0.2 to 1pmol of assembly products in a ratio between 1:1 and 1:5 of inserts to backbone was used for the Gibson assembly following the manufacturer’s instructions.

The assembled constructs were then cloned using XL-10 chemically competent cells (Agilent), and plasmid DNA was then purified using a commercial kit (Macherey-Nagel) followed by capillary sequencing. Capillary sequence files were trimmed using Benchling software and sequence alignments to confirm correct DNA fragment insertions were generated using algorithms such as ClustalOmega. Positive colonies were expanded in 200mL LB broth supplemented with 100ug/uL Ampicillin and incubated for 18h at 37°C overnight under shaking conditions at 200rpm. Plasmid DNA was then purified using a midi prep kit (Macherey-Nagel) in preparation for transfections. The cut pDC2-cam-coCas9-U6-yDHOODH was then dephosphorylated to avoid re-ligation in subsequent enzymatic steps. The gRNA sequences were pre-phosphorylated with adaptor sequences complementary to cut pDC2-cam-coCas9-U6-yDHOODH that were then used for the ligation step. Cloning was done using XL-10 chemically competent cells (Agilent) followed by capillary sequencing to identify positive clones from which plasmid DNA was purified in preparation for transfections using a midi prep kit (Macherey-Nagel).

### P. falciparum transfections

The purified targeting constructs were initially precipitated to generate a concentrated mixture that was also free from contaminating salts. To do this, a total of 60ug assembled pCC1 and 30ug assembled pDC2-gRNA plasmids for each target gene were pooled and mixed with 1/10 volume of 3M sodium acetate of the total plasmid volume. Samples were mixed by flicking the tube 4-5 times. 1mL of cold 100% Ethanol was then added to the sample and mixed by inverting 10 times followed by centrifugation at 15000rpm for 30min at 5°C. The supernatant was then discarded followed by addition of 1mL of cold 70% Ethanol and centrifugation at 15000rpm for 30min at 5°C. The supernatant was then discarded, and the sample further pelleted at 15000rpm for 30min at 5°C. The supernatant was then discarded, removing as much ethanol as possible without disturbing the pellet, and thereafter allowing the pellet to dry until it was translucent. The plasmid DNA was then resuspended in 50uL Cytomix solution ready for transfection.

Transfections were performed using ring-stage parasites. Cultures of *P. falciparum* NF54 parasite line at 4-5% parasitaemia were initially washed in warm 30 mL Cytomix per 2mL of packed iRBCs at 1500rpm for 5min and the supernatant discarded. A total of 300uL warm Cytomix was then added per 150uL of iRBC pellet. The resuspended DNA was then mixed with 450uL of the iRBC/Cytomix mixture and transferred to a 0.2cm Bio-Rad cuvette and thereafter electroporated using a BioRad electroporator (Gene Pulser Xcell, model number 1652660) using the following settings: Exponential mode, 310V Resistance: infinite, 950μF Cuvette: 2mm. After electrorotation, samples were then taken out of the cuvette and transferred to a 5mL culture 6-well plate, mixed with fresh RBCs at 2% hematocrit and incubated at 37°C ready for the next day.

### Post transfection parasite culture

The next steps after transfection were to maintain and keep the cultures on drug selection as needed until the desired mutants could be detected by microscopy. Positive selection was performed using 2.5nM WR99210 (Jacobus) and this was initially added to the cultures one day post the transfection. Smears were then made on day-2 post transfection to check for parasite death. Drug media was changed every day for the first six days post transfection followed by a 2-3-day cycle per week. Fresh uninfected RBCs were added every week post drug selection by discarding 1/3 of old culture and replacing this with an equal amount of fresh RBCs. Smears were performed twice a week until parasites appeared. The time taken for parasites to be detected under WR99210 selection varied from 8-21 days. Positive cultures were then frozen down until rounds of transfections had been completed, thus generating an arrayed library of resistant lines. Once the list of transfections had been completed, frozen lines were then thawed out and grown in 2.5nM WR99210 (Jacobus), followed by culture in a combination of both negative and positive selection using a mixture of 7mM 5-flurocytosine [5-FC] (Sigma) and 2.5nM WR99210 (Jacobus) respectively for three cycles. Parasite were grown for at least three cycles in complete medium supplemented with 7mM 5-flurocytosine [5-FC] and 2.5nM WR99210 to select ideally for double-crossover recombinants. Resistant parasites were then expanded out and pellets directly frozen at either –20°C for genotyping or in glycerolyte stocks at –80°C for future recovery and use.

### Genotyping of deletion parasites

Transgenic parasites were genotyped to confirm the desired gene disruptions. Capillary sequencing using a conserved primer present in the cassette of all the targeting constructs was performed to verify the identity of the barcode module that the parasites were carrying. PCRs were also performed to assess gene disruptions by agarose gel electrophoresis. Primers were designed to exclusively amplify either the wild-type or the transgenic locus and therefore distinguish modified and unmodified loci. A complete list of all primers used both in plasmid construction and genotyping is shown in **Supplementary Table 3**.

### BarSeq

Barcode Sequencing (BarSeq) was used in multiple assays to test relative growth rates, DNA content and cell cycle dynamics of deletion strains. In these experiments, genomic DNA was extracted from parasite DNA pellets using the Qiagen DNeasy Blood & Tissue Kit (Qiagen), following the manufacturer’s instructions. A nested PCR was performed to generate the BarSeq sequencing libraries.

For the first PCR which attaches the adaptor overhangs, each reaction mix had a total of 25uL, containing, 0.2uM of the forward and reverse primers (sequences), distilled water, 2uL of DNA template, and 1X of 12.5uL CloneAmp containing both the DNA polymerase enzyme and dNTPs. Samples were amplified at an initial denaturation temperature of 95°C for 5min, followed by 30 cycles of denaturation at 95°C for 30s, then annealing at 54°C for 30 s, and extension at 72°C for 3s. Samples were finally extended for 5min at 68°C.

For the second PCR which attaches the index barcodes needed for cluster generation and demultiplexing, samples were amplified using index primers from Illumina that are unique to each experimental condition. Each reaction volume contained a total 0.2uM of both the forward and reverse index primers, 2uL of neat amplicon product from the first PCR, 12.5uL of CloneAmp containing both the DNA polymerase enzyme and dNTPs, and water to make up a total volume of 25uL. Thermocycling conditions were set at an initial temperature denaturation of 95°C for 2min, followed by 12 cycles of denaturation at 95°C for 30s, then annealing at 68°C for 15s, and extension at 68°C for 8s. Samples were finally extended for 5min at 68°C.

Prior to next generation sequencing, libraries were initially cleaned using a Qiagen MiniElute UF PCR purification kit (Qiagen) as per the manufacturer’s instructions. The concentration of each of the purified samples was initially measured using the Quant-iT PicoGreen dsDNA reagent (Thermo Fisher Scientific) or Qubit. 125ng of each sample was then pooled and re-measured using the Qubit fluorometric broad sensitivity quantitation kit (Thermo Fisher Scientific). The DNA pool was then diluted to a concentration of 4nM and sequenced at the Wellcome Sanger. Libraries were run using a Miseq Reagent Kit v2 Paired-End Reads 2 x 150bp with the cluster generation and sequencing all performed using a Miseq system, which generated 24-30 million reads

### Growth measurements using arrayed and pooled approaches

For the arrayed approach, the lines were divided into four subgroups of 22-27 lines each, which were then assayed sequentially to each other. This approach was the only one that was technically feasible, and took a period of three months, but had the risk of increasing variation in growth phenotypes due to batch effects. To allow for comparison between experiments, eight controls – PfRH3 (variant 1), PfRH3 (variant 2), PfPARE, PfRNA binding (PF3D7_1241400), PfASP (PF3D7_0405900), PfPARE [Dd2], PfMyoB (PF3D7_0503600) and PfEBA165, were included as much as possible in each of the four subgroups. To allow growth to be compared between groups and both the arrayed and pooled approaches, growth was expressed relative to an average of the two PfRH3 control lines and these were also used to assign qualitative growth phenotypes using confidence intervals.

Growth assays for the arrayed approach were performed starting with either asynchronous or synchronous cultures. In the latter, parasites were enriched using LD MACS columns. To do this, the LD MACS columns were placed on a magnetic stand and equilibrated with 3mL RPMI, and thereafter loaded with 4mL of parasites at 50% haematocrit. Once the flow through with iRBCs was complete, the columns were washed 2X using 4mL RPMI and thereafter eluted off the magnet in 4mL RPMI. The eluate at this point was now enriched with late-stage parasites which were then allowed to re-invade fresh uninfected RBCs. Experiments were then seeded the following day, with parasite cultures that were predominantly (>90%) ring stages. The asynchronous stages were seeded with variable proportions of rings, trophozoites and schizont stage parasites.

Mixed pools of parasite deletion lines were maintained in either static or shaking culture at 60rpm on an INFORS HT bench-top shaker and cultured in the same incubator. A simple index to formally assess growth phenotypes between shaking and static conditions for each deletion line was generated and referred to as the shaking vs static index (SSI). The RH3 controls were used as baseline and phenotypes were scored as low if the mean SSI_GR_ estimate of the deletion lines was lower than the PfRH3 controls and with non-overlapping confidence intervals, non-significant if the deletion lines had overlapping confidence intervals with the PfRH3 controls or high if the mean SSI_GR_ estimate of the deletion lines was higher than the PfRH3 controls and with non-overlapping confidence intervals.

### DNA content measurements

Parasites at mid-schizont stages were treated with Compound 2 drug to block egress of mature segmented schizont stages (Donald, Zhong et al. 2006), stained with 4uM Hoechst 33342 and staining intensity acquired on a Fortessa (BD Biosciences) flow cytometer. The average total DNA content per schizont from the arrayed experiments was determined by assessing the difference in relative fluorescence units of schizont stage parasites normalized to the fluorescence units in ring stage parasites, which contain only one copy of the *P. falciparum* genome. Experiments were performed in four subgroups. Additional parameters such as kurtosis and skewness were also determined. A total of six controls including PARE [Dd2], PfRH3 (variant 1), PfRH3 (variant 2), PfEBA165, PfPARE and MyoB were included in each of the subgroups. The two PfRH3 lines PfRH3 (variant 1) and PfRH3 (variant 2) were used to determine the boundaries of the confidence intervals for phenotype assignment. The DNA content of the deletion lines was scored as either low (if the mean DNA content was lower that the controls and had non-overlapping confidence intervals to the PfRH3 controls), or non-significant (if the confidence intervals of the mean DNA content overlapped with the PfRH3 controls) or high (if the mean DNA content was higher than the PfRH3 controls and with non-overlapping confidence intervals to the PfRH3 controls).

For BarSeq assessment of schizont DNA content, mixed pools of synchronous parasite cultures at mid-schizont stages were incubated in 1.5uM Compound 2 for 8 hours, fixed in 25% Methanol for 15 min, washed with 1X PBS, stained with 4uM Hoechst 33342, and then sorted into five operational gates (shown in **Supplementary Figure 11**) using the BD influx cell sorter (BD Biosciences). The total number of cells sorted ranged from 2000 – 150000. These volumes were then spun down and frozen. A volume of 2uL of this were then used for the BarSeq experiments as detailed in the methods to generate reads. A frequency distribution table was used to calculate the mean barcode content and confidence intervals. Significance in the mean barcode content was determined by comparing each deletion line to PfRH3 (variants 1 and 2). The barcode content of the deletion lines was scored as either low (if the mean barcode content was lower that the controls and had non-overlapping confidence intervals to the PfRH3 controls), or non-significant (if the confidence intervals of the mean barcode content overlapped with the PfRH3 controls) or high (if the mean barcode content was higher than the PfRH3 controls and with non-overlapping confidence intervals to the PfRH3 controls).

### IDC progression

Synchronous parasites at schizont stages were pooled and allowed to invade uninfected RBCs on a shaker at 60rpm. After 3-6 hours, the pooled cultures were treated with 5% Sorbitol to remove non-egressed schizonts. For the proof of principle experiment, samples were sorted at intervals of 6h over a 48h time course using the BD influx cell sorter (BD Biosciences). At each time of sampling, 125uL of a 2% culture was fixed in 25% Methanol for 15 min, washed with 1X PBS and then stained in 2mL mixture of 4uM Hoechst 33342/ 0.1ng/mL Thiazole Orange until parasites were ready for acquisition. The total number of cells sorted in each gate ranged from 2000 – 150000 from multiple gates as shown in the results and density was dependent on the stage of the cycle. These volumes were then spun down and frozen down. A volume of 2uL of this were then used to extract gDNA for the BarSeq experiments.

Two methods were used to measure the transition rates of parasites through the gates – a ‘before and after’ weighted approach, and also a compartmental model to simulate the transition of reads between gates. Each compartment records the proportion of sequencing reads observed at time *t* in that gate. We modelled independent transition rates 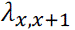 for each pair of sequential gates.

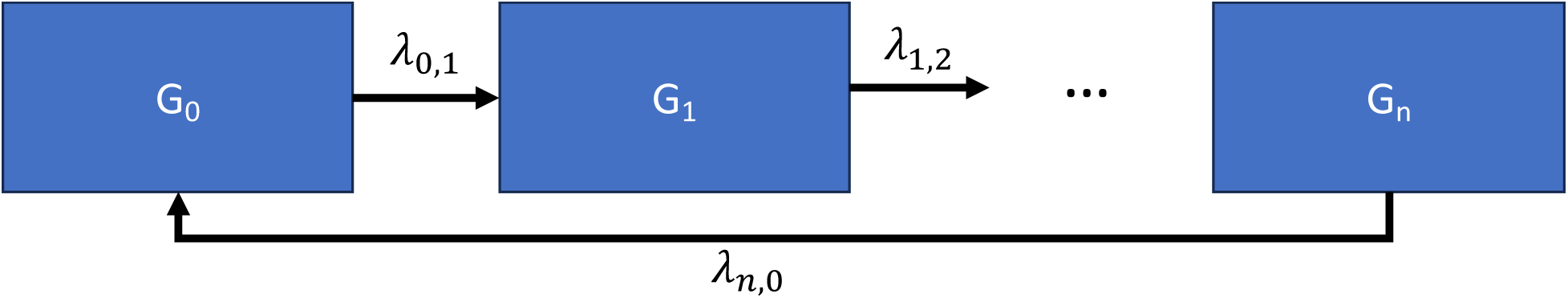

Basic structure of the compartmental model.

For each gate, the change in read proportion is modelled as:

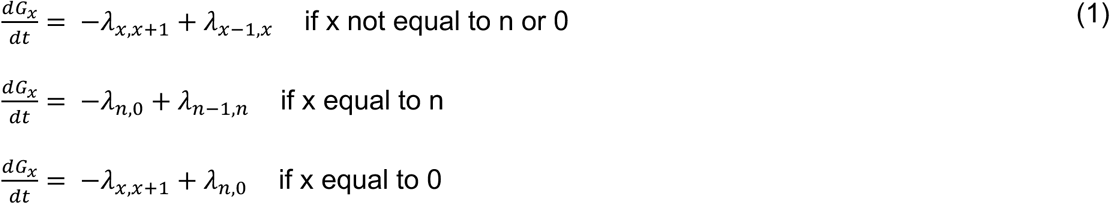

where 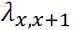 is the transition rate from the current gate to the next gate, 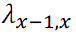 is the current gate to the next gate, and *dt* is measured in hours. Note that the model assumes that the transition rates are constant through time. It furthermore assumes that all the sequencing reads in a given compartment will eventually transition to the next compartment.

The transition rates of the compartmental model were independently calibrated to each of the genes examined. These rates were calibrated by minimizing the mean square error between the proportion of reads in each gate collected at hour *t* and the simulated results at time *t*.

For the bigger pools, we used a weighted ‘before and after’ approach to assess the transition rates of parasites through the IDC.

### Modelling the effect of changes in DNA content and cell cycle progression on growth rates

The combination of DNA content and IDC progression were selected as the independent variables (X), and parasite growth rates chosen as the dependent variable (y). The dataset was then divided into training and testing sets using an 80/20 split through the train_test_split function from scikitlearn. To ensure that the features were on a similar scale, particularly important for the models used, the features were standardized using StandardScaler. For the Support Vector Regression (SVR) model, the features DNA content and IDC progression were standardized using StandardScaler to ensure that each feature contributed equally to the model. The SVR model was implemented using the SVR class from scikit-learn, with a polynomial kernel (kernel=’poly’), a regularization parameter C=100, and an epsilon value of 0.1. The polynomial kernel was selected to capture the non-linear relationships between the features and the target variable. The model was trained on the standardized training data, and predictions were made on the test set. The performance of the model was evaluated using the Mean Squared Error (MSE) metric, which resulted in an MSE of 0.016. A scatter plot was generated to visualize the relationship between the true and predicted values. Specific values were also defined and standardized for prediction, demonstrating the model’s prediction capabilities for unseen data points. A Dense Neural Network (DNN) was constructed using TensorFlow’s Keras API for the task of regression. The features DNA content and IDC progression were standardized prior to training to ensure consistent input to the model. The network architecture consisted of three fully connected layers: the first layer with 64 neurons and ReLU activation, the second layer with 32 neurons and ReLU activation, and a final output layer with a single neuron to predict the continuous target outcome variable. The model was compiled using the Adam optimizer and the Mean Squared Error (MSE) loss function. Training was conducted over 20 epochs with a batch size of 32, and a validation split of 20% was used to monitor the model’s performance on unseen data during training. The model achieved a test MSE of 0.059. Additionally, a plot of the training and validation loss across epochs was generated to assess the model’s learning process. The trained DNN was used to make predictions on both the test set and specific predefined values, which were standardized similarly to the training data, showcasing its ability to generalize to new inputs. For the Random Forest Regression model, the RandomForestRegressor from scikit-learn was employed with 100 trees (n_estimators=100). The model was trained on the standardized training data, and predictions were made on the test set. Model performance was assessed using the Mean Squared Error (MSE) metric, yielding an MSE of 0.028. The relationship between the true and predicted values was visualized with a scatter plot. Additionally, specific values were defined for prediction purposes, standardized, and passed through the trained model to obtain predicted outcomes, demonstrating the model’s ability to generalize to new data points.

### Data analysis

Fastq files were generated from the illumina sequencing pipelines and contained raw barcode counts. An in-house script in Perl was used to generate read counts in excel files that could then be manipulated using different statistical methods and software. Scripts in R software were generated to analyse both the arrayed and BarSeq data. A range of statistical approaches were used nd details of these are discussed in the results sections.

