## Supplementary Material for "Application of barcode sequencing to increase the throughput and complexity of *Plasmodium falciparum* genetic screening"

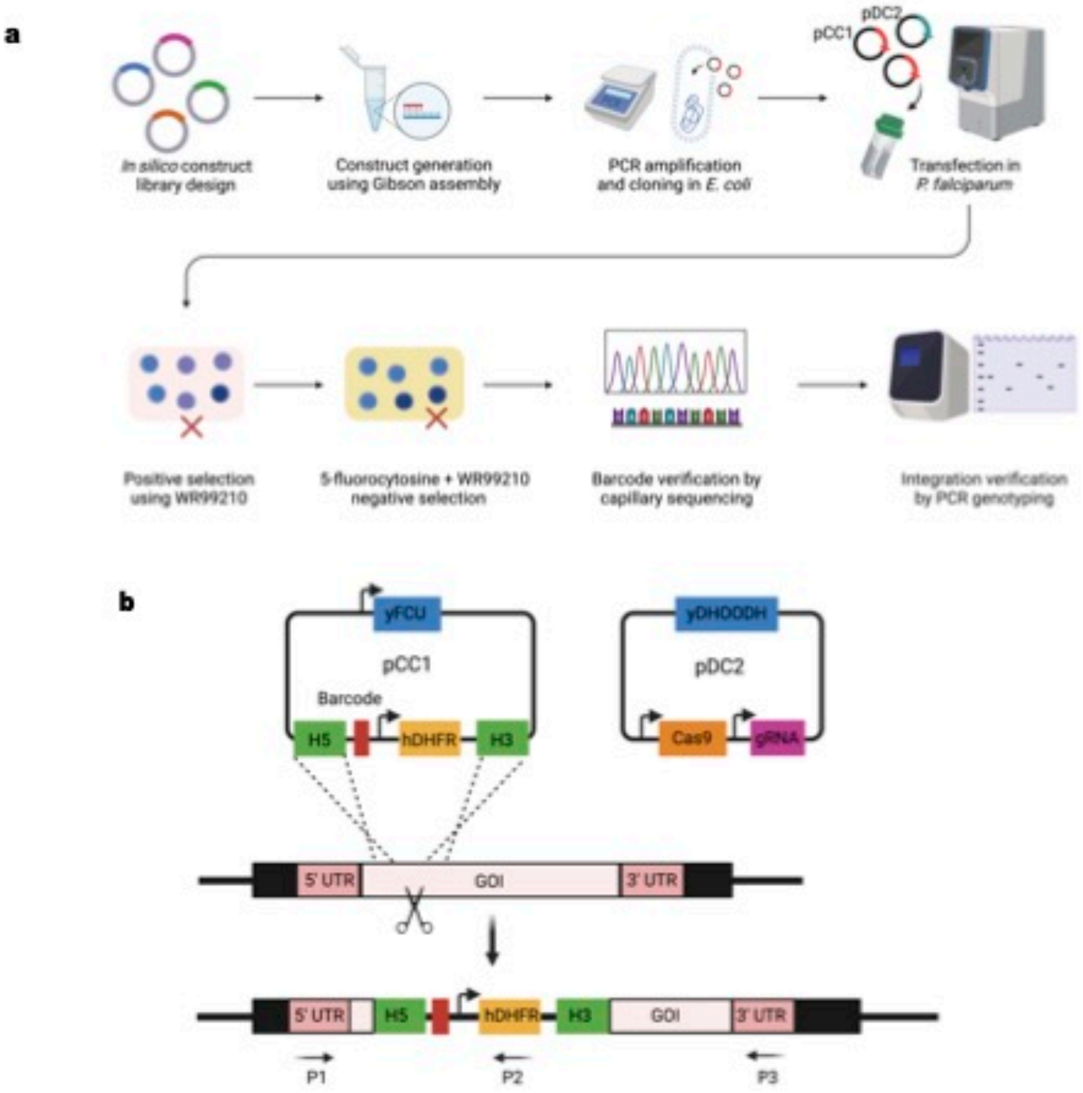


**Supplementary Figure 1. Overview of the CRISPR/Cas9 pipeline used to generate barcoded *P. falciparum* deletion lines.** (a) A Library of gene deletion constructs targeting a selection of loci were initially designed *in silico* and constructs assembled experimentally thereafter using the Gibson assembly method (Gibson, Young et al. 2009). Plasmid constructs were then transfected into *P. falciparum* NF54 wild-type ring stage parasites by electroporation. A two-plasmid approach was used as shown in (b), with a combination of pCC1 (carrying the repair template, hDHFR positive marker and yFCU negative selection marker) and pDC2 (carrying both the gRNA and

Cas9 enzyme). Transfected parasite cultures were then selected initially on WR99210 drug pressure until parasite proliferation could be detected by light microscopy. This was then followed by treatment of lines with a combination of WR99210/ 5-fluorocytosine (5FC). Parasites that were resistant to the combination of WR99210/ 5-fluorocytosine (5FC) were then genotyped using capillary sequencing to verify presence of the barcode insert as well as conventional PCR for evidence of locus deletion using a combination of primers P1, P2, P3 as shown in panel (c). Created with BioRender.com

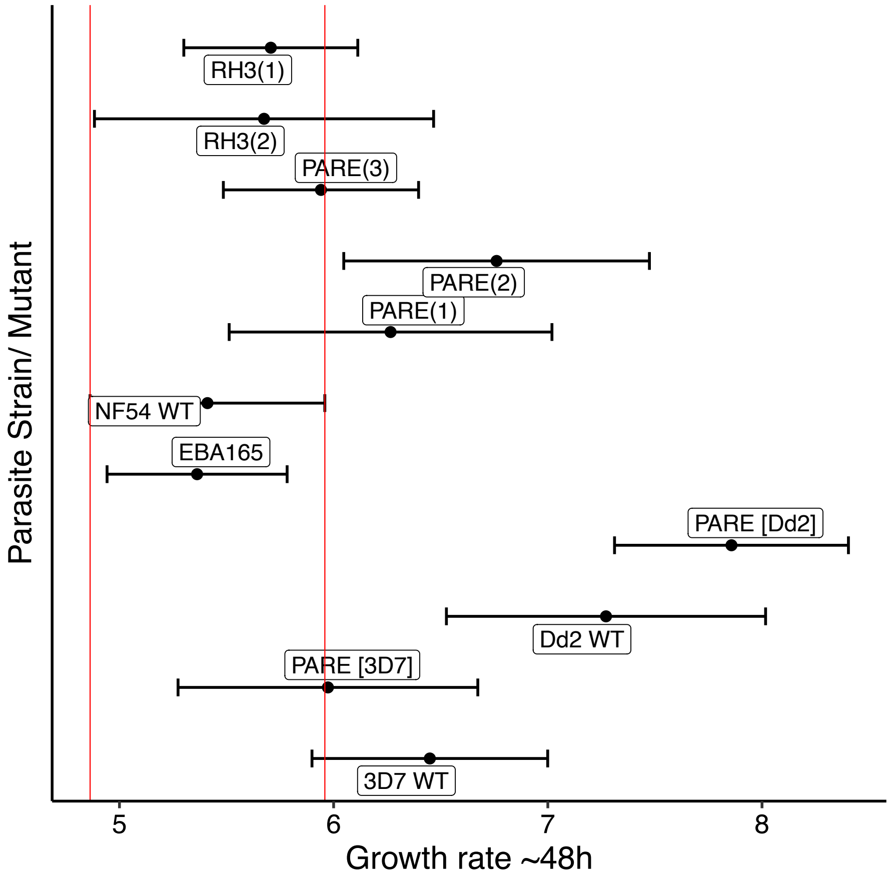


**Supplementary Figure 2. Growth rates of control deletion lines.** Comparisons were made between wild type parasite strains (NF54, 3D7, Dd2) and deletion lines within background of either

NF54 (RH3 variant 1, RH3 variant 2, PARE, EBA165) , 3D7 (PARE) or Dd2 (PARE). Boundaries of NF54 wild type confidence intervals were used to assign significance (red).

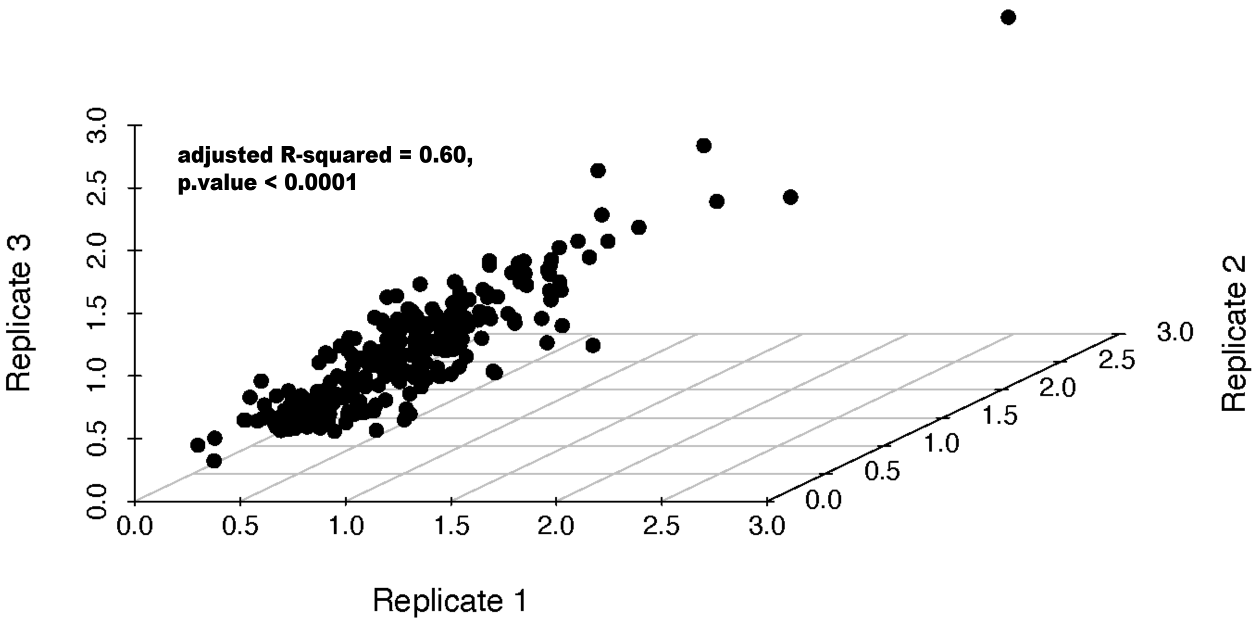


**Supplementary Figure 3. Correlation of replicates from pooled BarSeq assay in static culture.** Replicates of growth rates per 48h were compared for each deletion line in pooled cultures under static conditions.

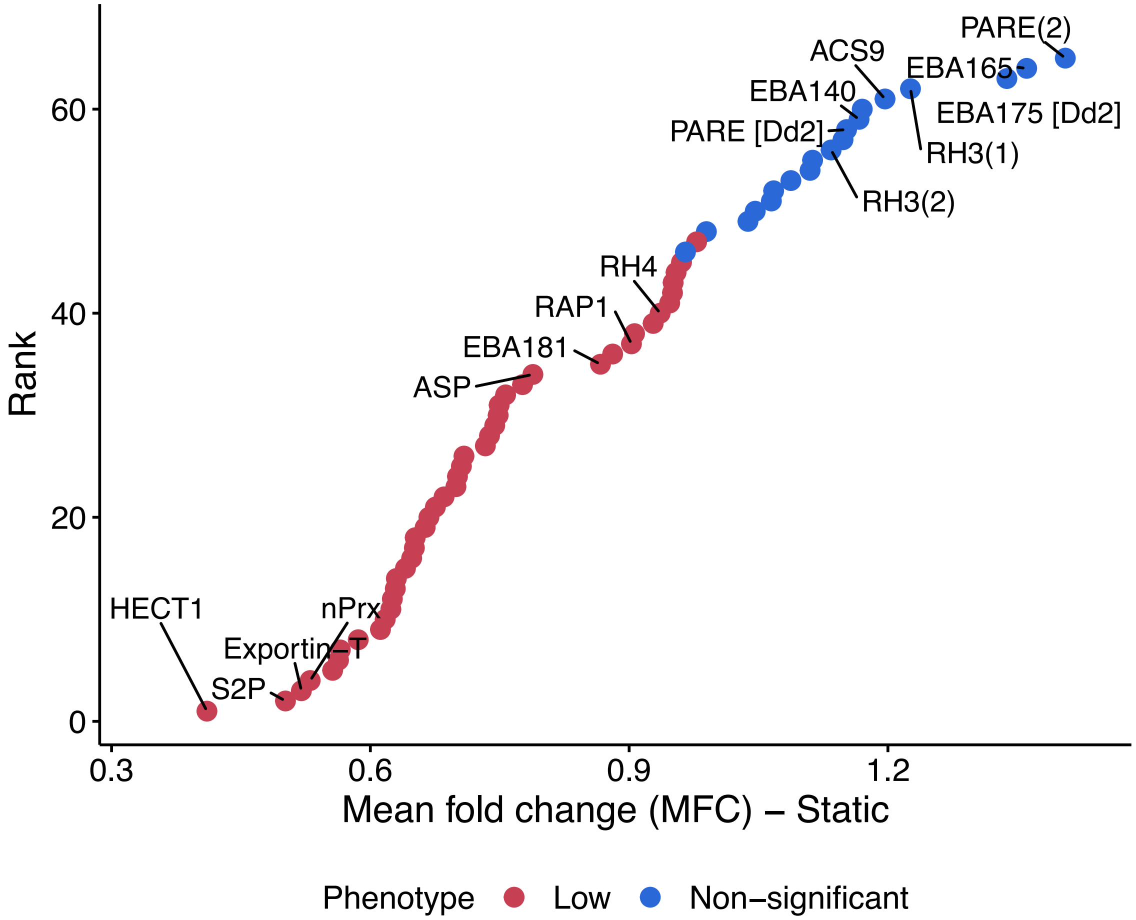


**Supplementary Figure 4. Ranked order of deletion lines in pooled BarSeq assay in static culture.** Relative growth rates of deletion lines in static culture ordered by rank in RGR estimate

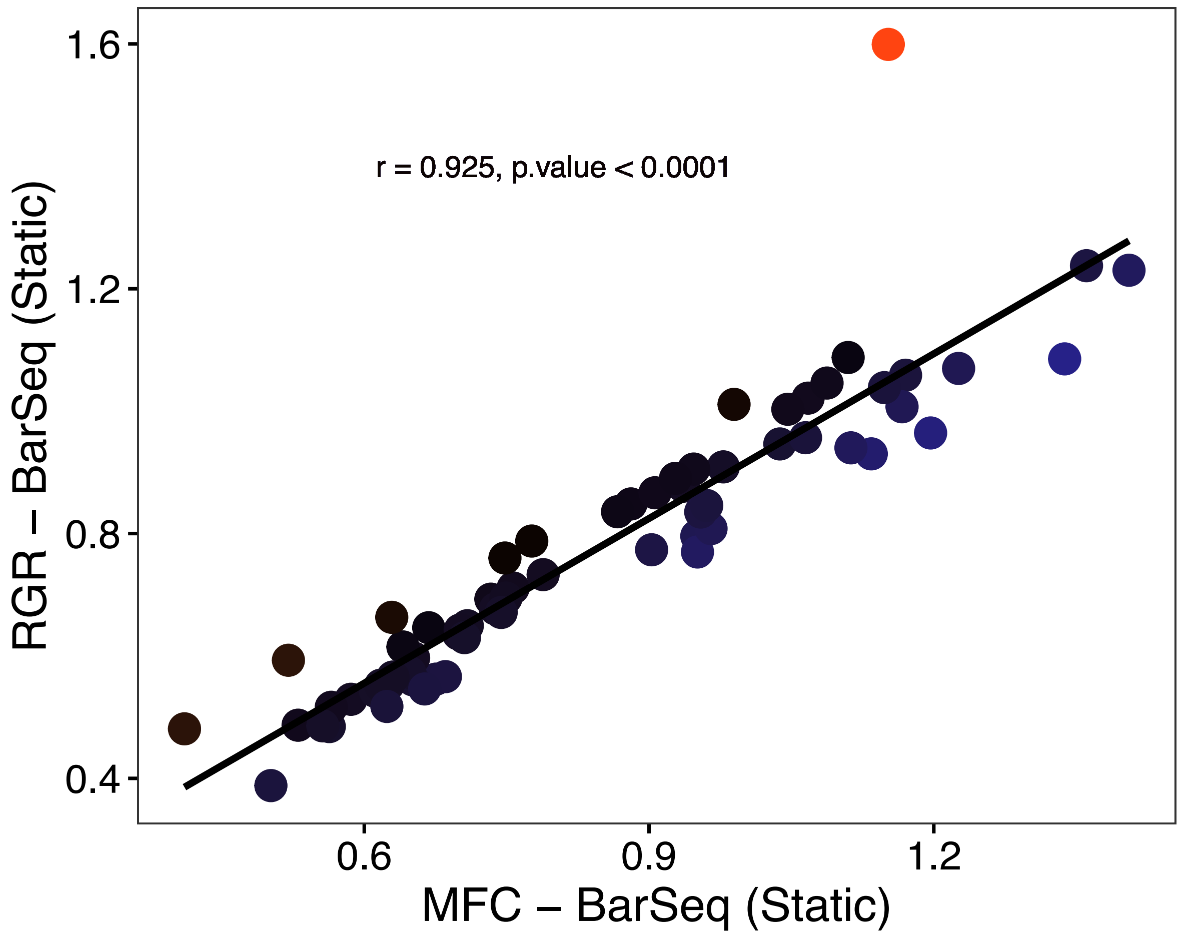


**Supplementary Figure 5. Correlation between methods for assessing growth rates in mixed pools in static culture**. Growth rates from BarSeq experiments were estimated using either relative growth rates and the RH3 variants (PfRH3 variant1 and PfRH3 variant 2) as controls or mean fold changes (MFC) that are independent of controls.

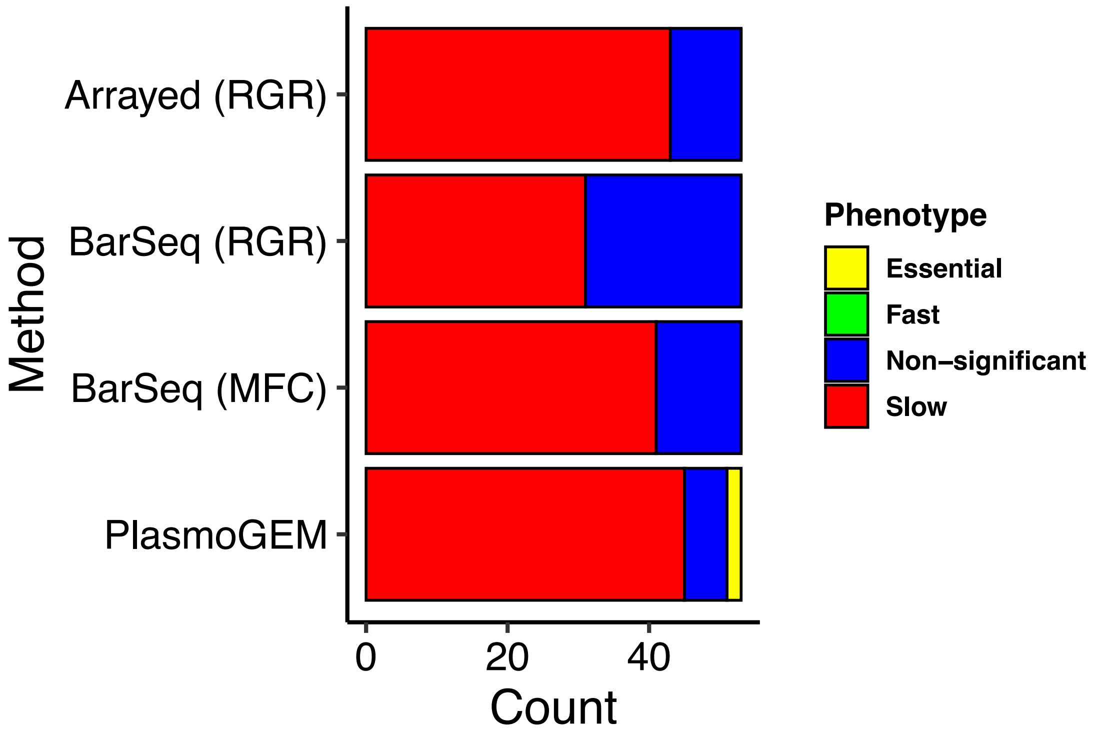


**Supplementary Figure 6.** Comparison of growth fitness phenotype assignments using the arrayed and mixed pool BarSeq approaches with published *Plasmo*GEM and *Piggy*Bac large-scale screens. Analyses were restricted to genes that were orthologous in *P. berghei*.

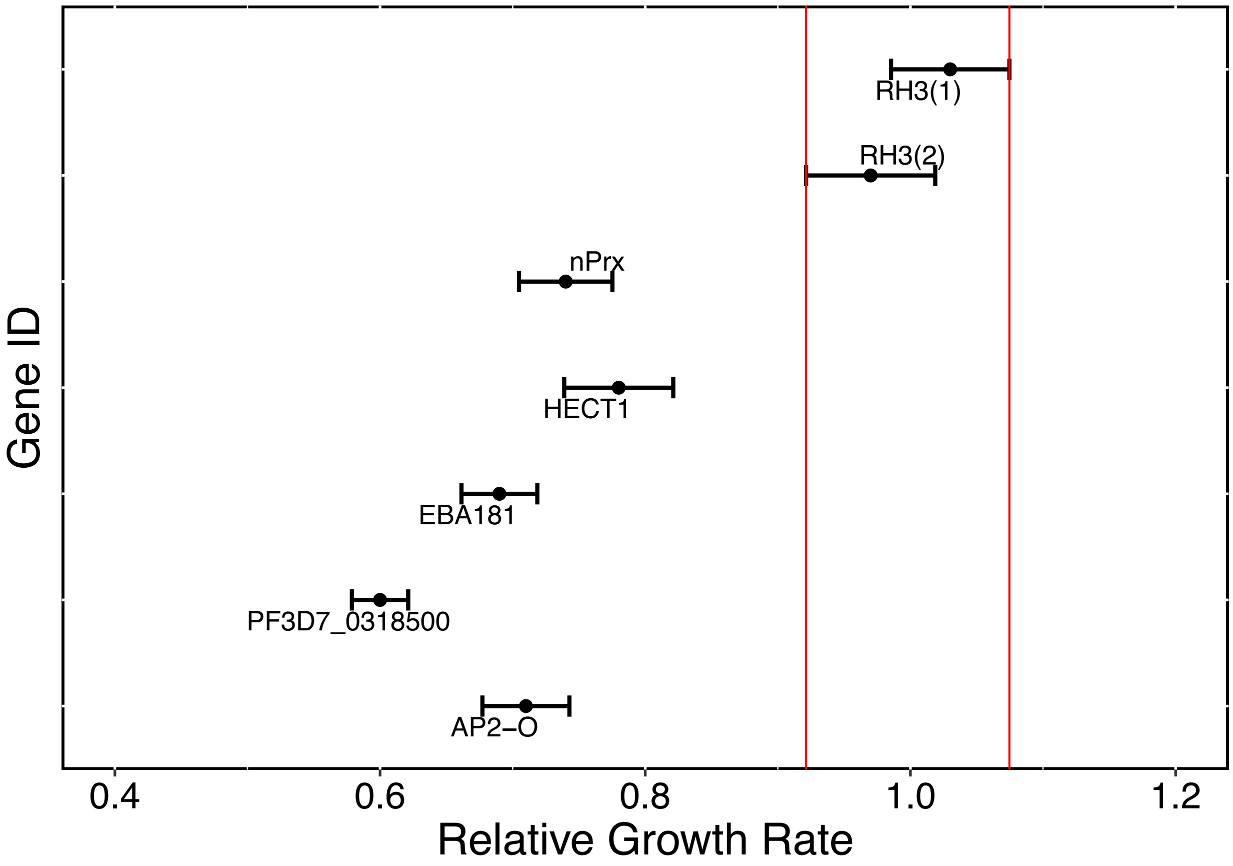


**Supplementary Figure 7. Growth rates of independent repeat deletion lines**. New constructs from those in the initial CRISPR/Cas9 pipeline were generated using Gibson assembly targeting three loci (HECT1, nPrx, and PF3D7_0318500) and transfected into NF54. Following drug selection and genotyping to confirm locus deletion and barcode insertion, growth assays were performed in an arrayed format for these new deletions alongside additional controls including PfRH3 (variant 1), PfRH3 (variant 2) as well as lines (PfEBA181, PfAP2-O) that had slow growth rates in the initial growth screen. Boundaries of confidence intervals for the two PfRH3 variants (red lines) were used to determine significance.

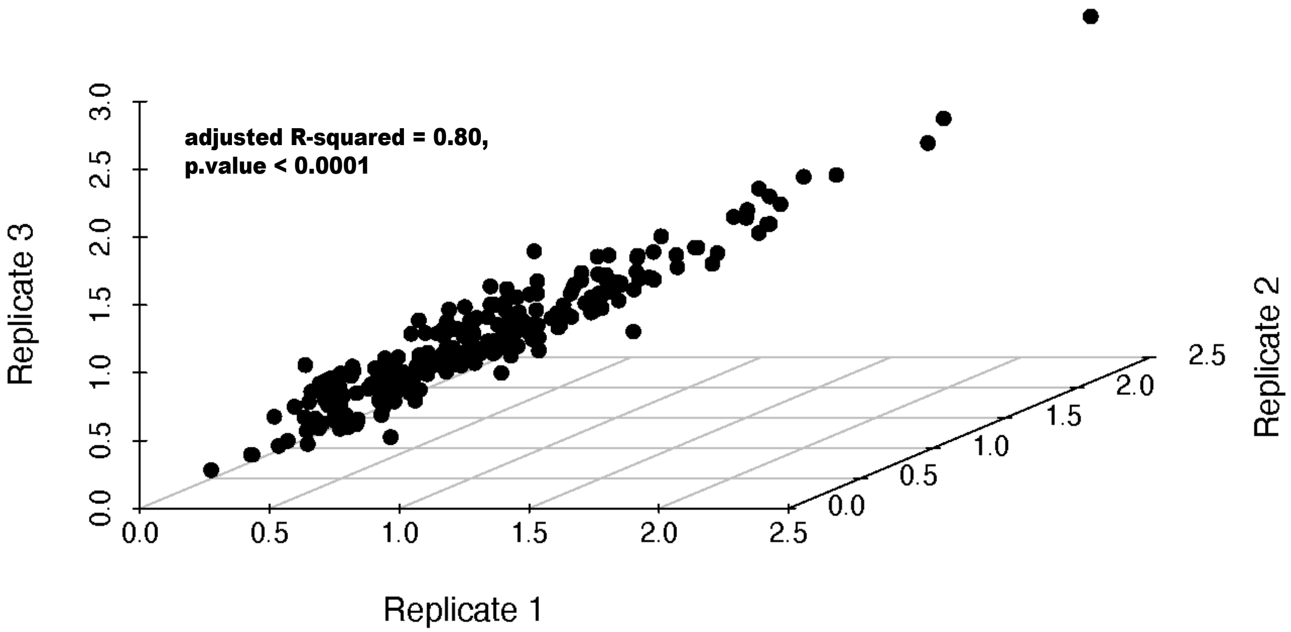


**Supplementary Figure 8. Correlation of replicates from pooled BarSeq assay in shaking culture.** Replicates of growth rates per 48h were compared for each deletion line in pooled cultures under shaking conditions.

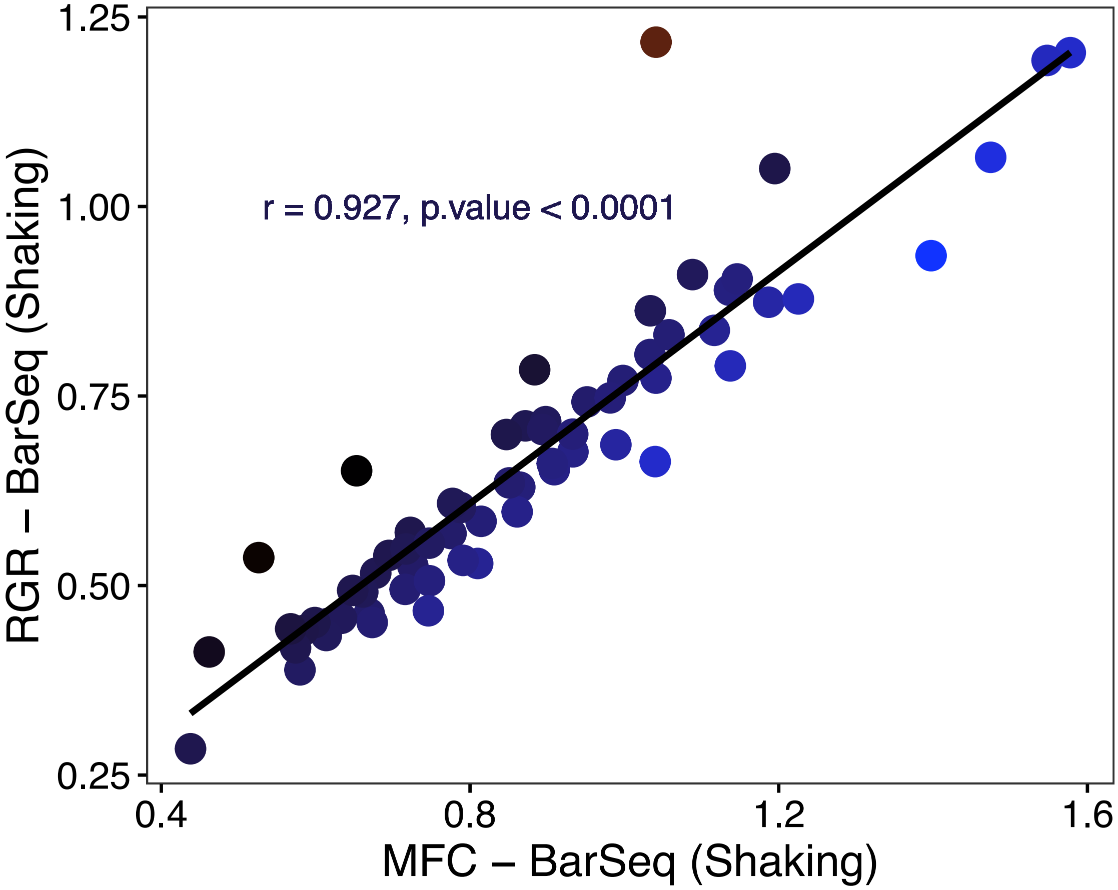


**Supplementary Figure 9. Correlation between methods for assessing growth rates in mixed pools in shaking culture**. Growth rates from BarSeq experiments were estimated using either relative growth rates and the RH3 variants (PfRH3 variant1 and PfRH3 variant 2) as controls or mean fold changes (MFC) that are independent of controls.

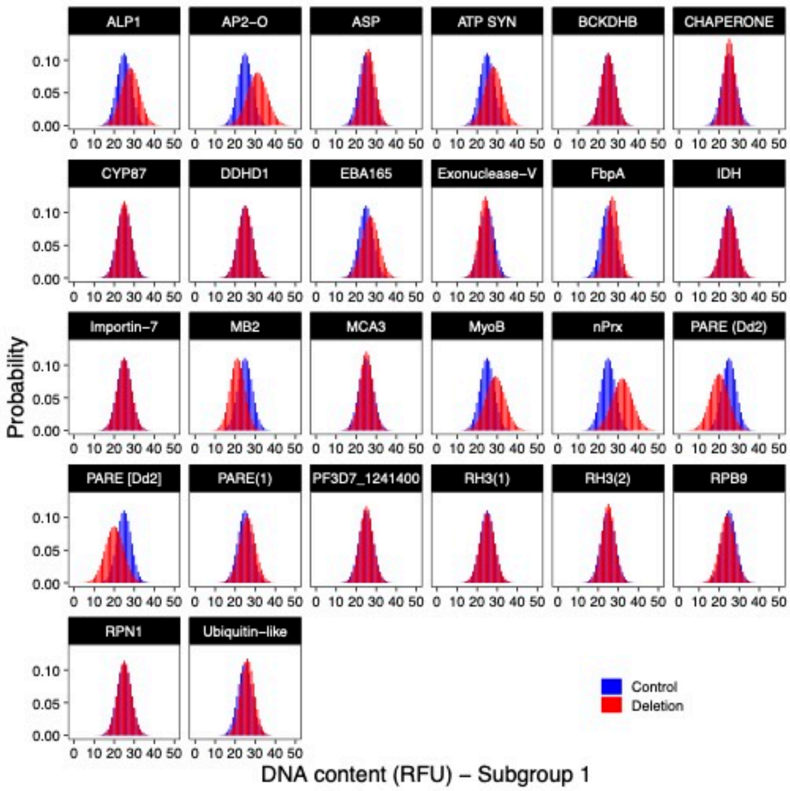
ALP1
AP2−O
ASP
ATP SYN
BCKDHB
CHAPERONE
CYP87
DDHD1
EBA165
Exonuclease−V
FbpA
IDH
Importin−7
MB2
MCA3
MyoB
nPrx
PARE (Dd2)
PARE [Dd2]
PARE(1)
PF3D7_1241400
RH3(1)
RH3(2)
RPB9
RPN1
Ubiquitin−like
Probability
Control
Deletion
DNA content (RFU) − Subgroup 1

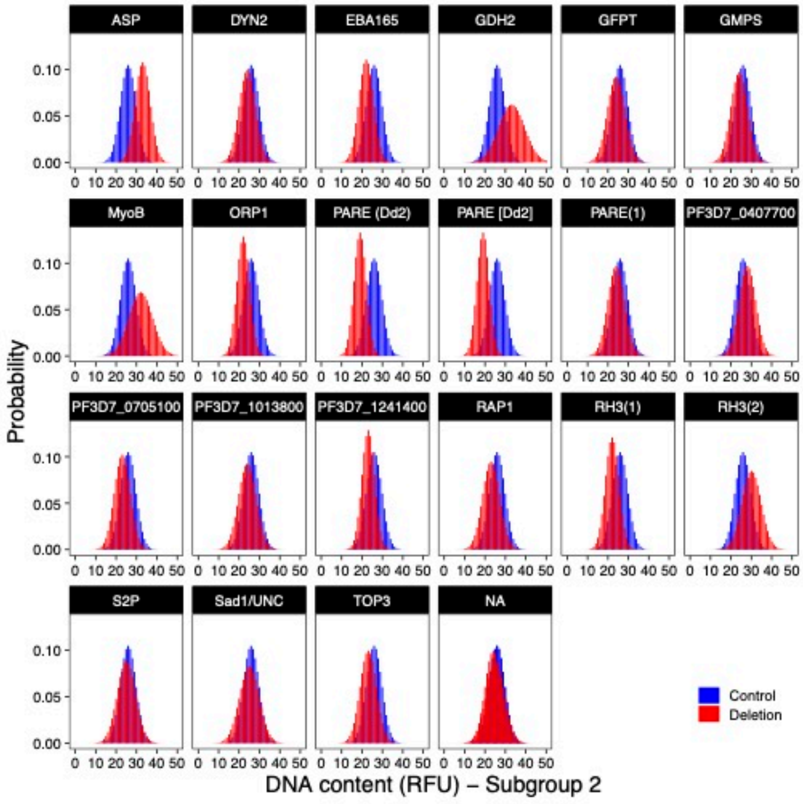
ASP
DYN2
EBA165
GDH2
GFPT
GMPS
MyoB
ORP1
PARE (Dd2)
PARE [Dd2]
PARE(1)
PF3D7_0407700
PF3D7_0705100
PF3D7_1013800
PF3D7_1241400
RAP1
RH3(1)
RH3(2)
S2P
Sad1/UNC
TOP3
NA
Probability
Control
Deletion
DNA content (RFU) − Subgroup 2

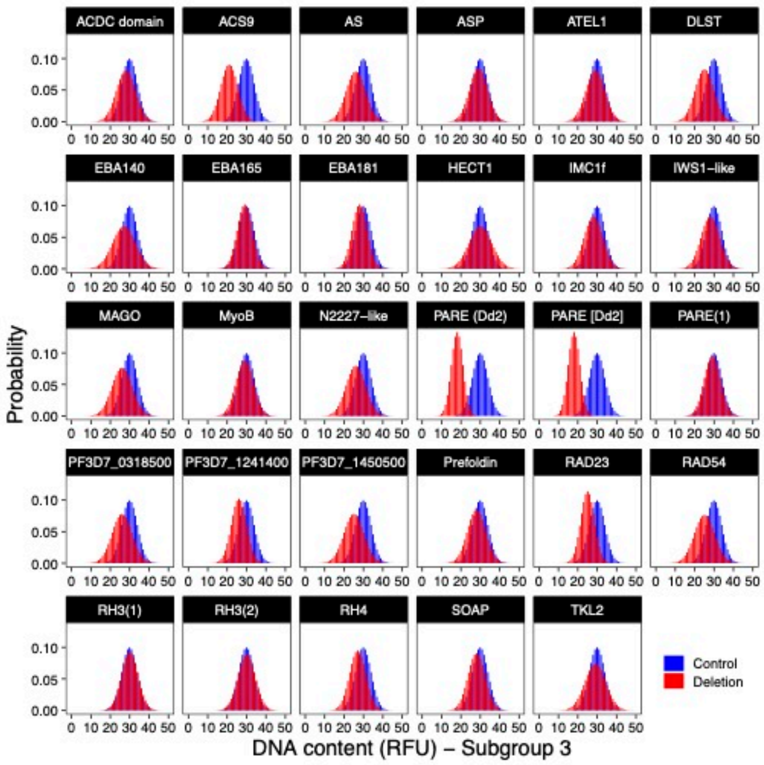


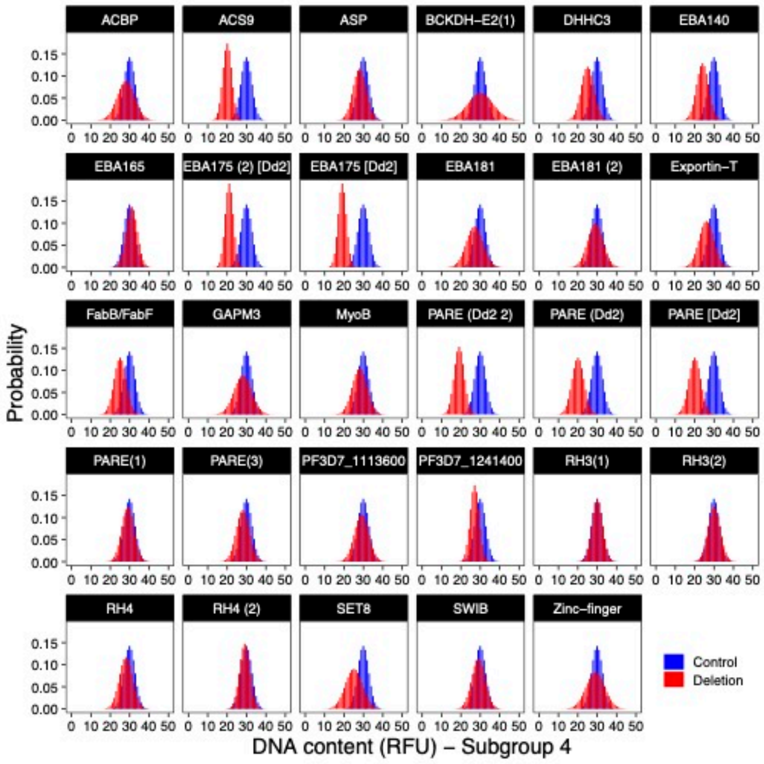


**Supplementary Figure 10. Distribution of DNA content of deletion lines using the arrayed approach.**

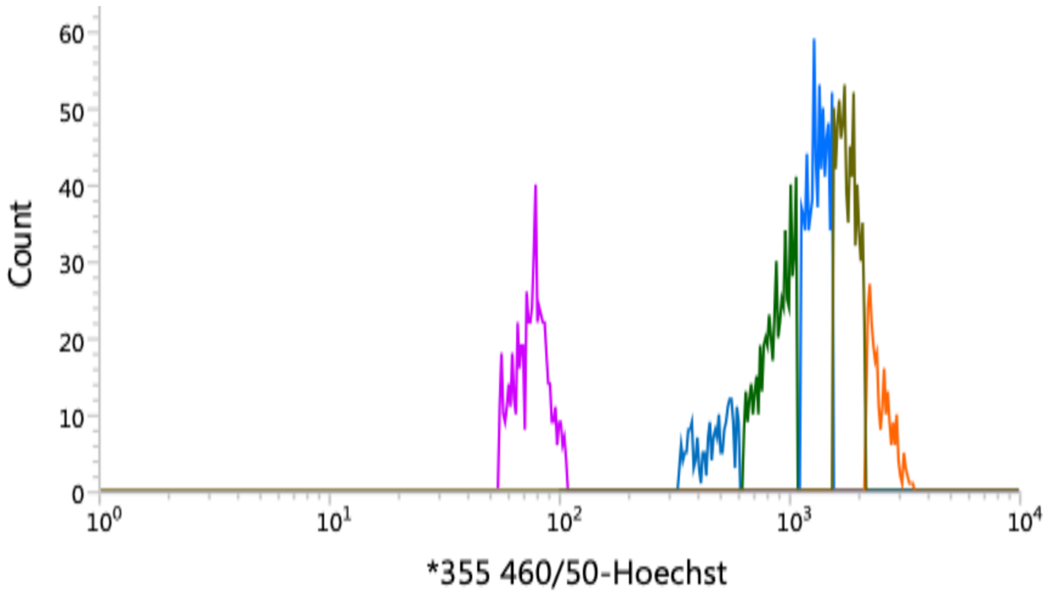


**Supplementary Figure 11 – Distribution of DNA signal in mixed pools.** Synchronous mixed pools of deletion lines were cultured and late stage schizonts were arrested just before egress using the Protein Kinase G inhibitor, Compound 2. The DNA content of these parasite lines follows a normal distribution that was then sub-divided into quintiles that were then used to sort separately parasite populations within each of the windows.

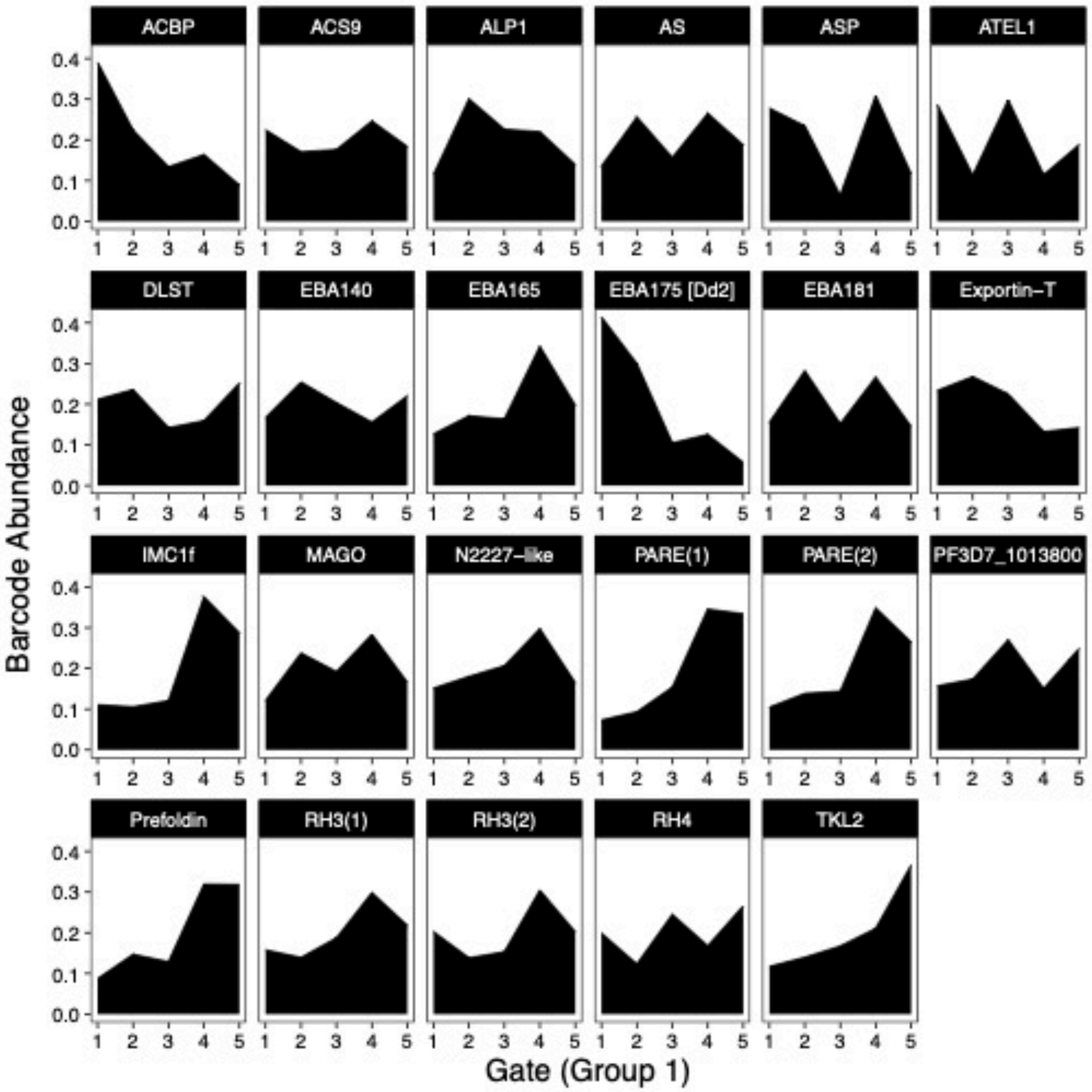

ACBP
ACS9
ALP1
AS
ASP
ATEL1
DLST
EBA140
EBA165
EBA175 [Dd2]
EBA181
Exportin−T
IMC1f
MAGO
N2227−like
PARE(1)
PARE(2)
PF3D7_1013800
Prefoldin
RH3(1)
RH3(2)
RH4
TKL2
Barcode Abundance
Gate (Group 1)


110

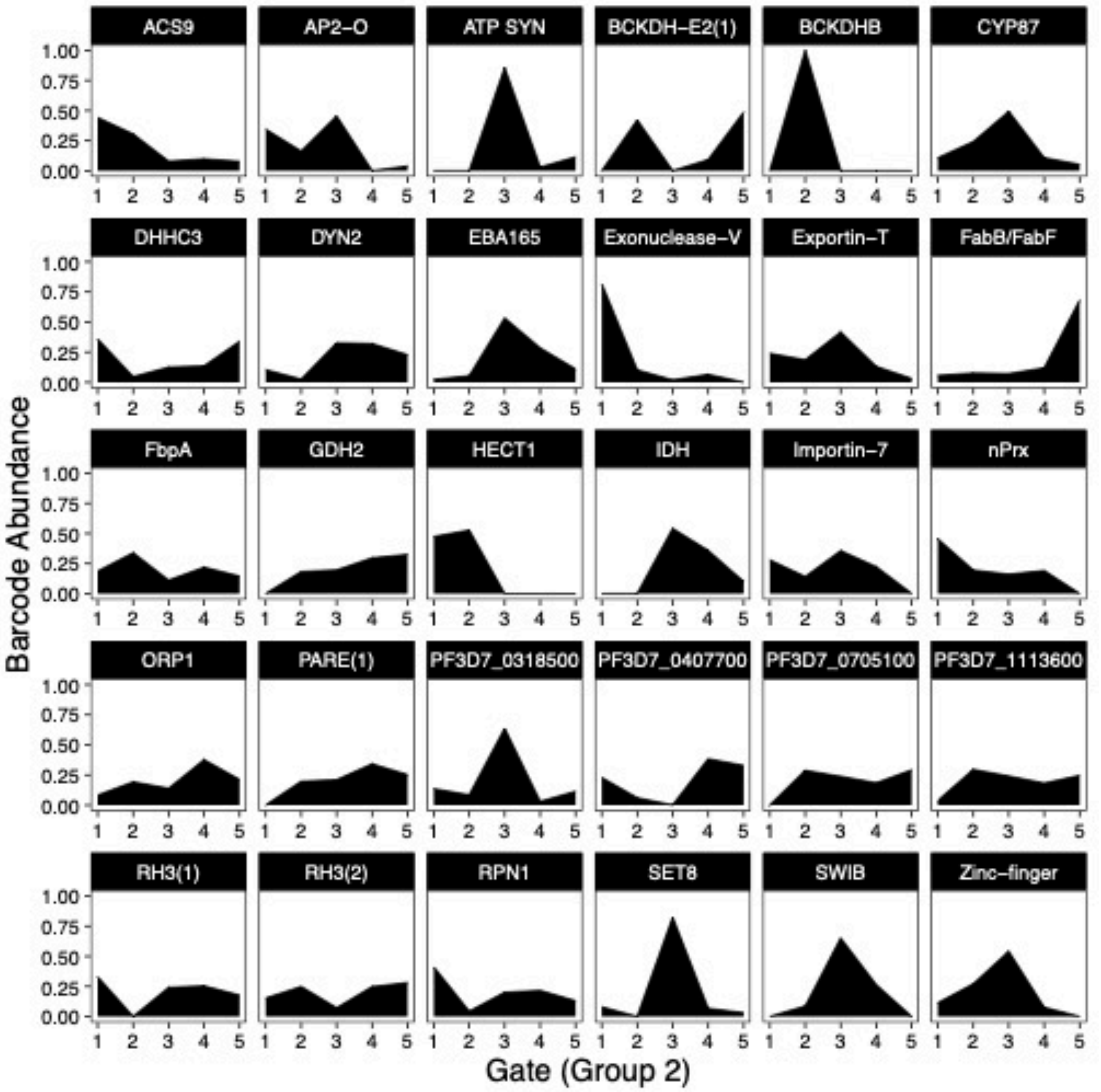


**Supplementary Figure 12. Distribution of barcode content of deletion lines from pooled culture**

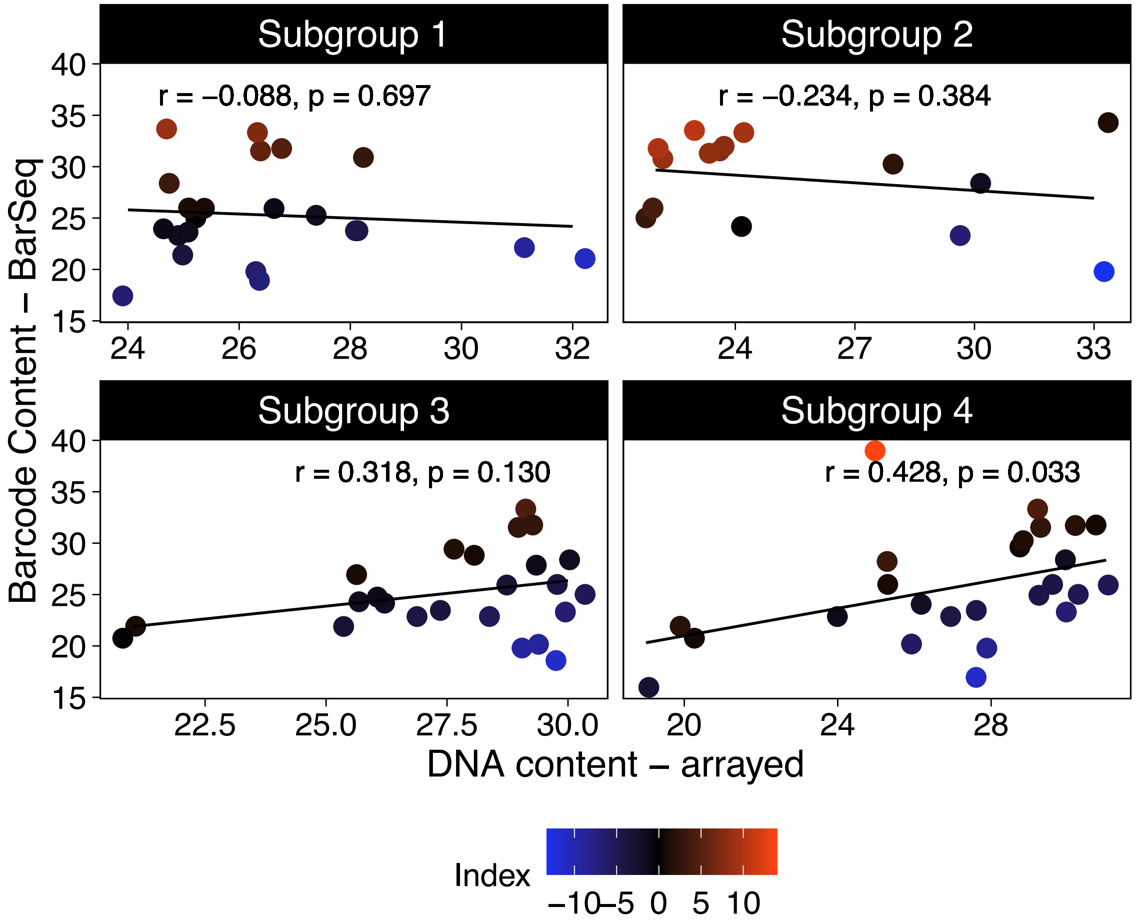


**Supplementary Figure 13. Comparison of arrayed and mixed pool approaches**

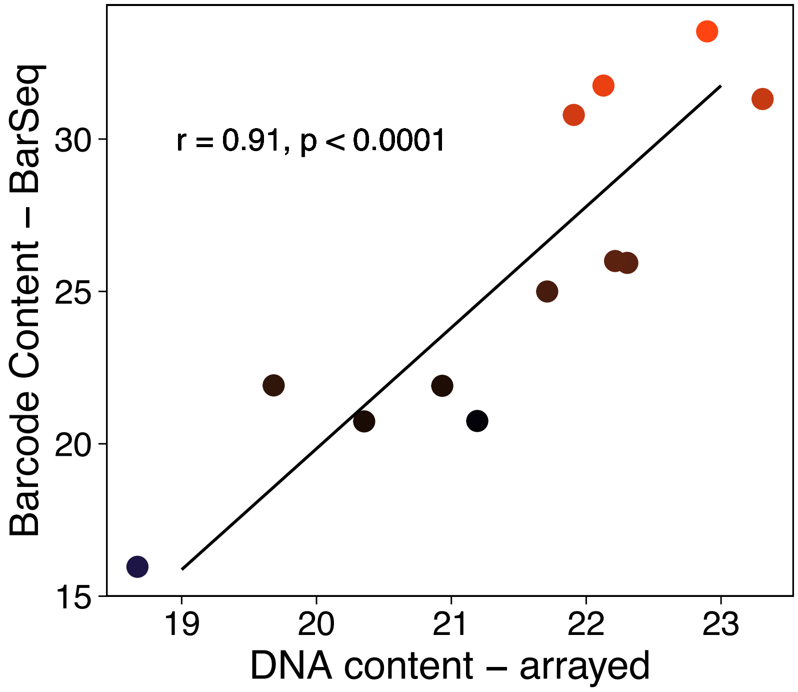


**Supplementary Figure 14. Comparison of arrayed and mixed pool approaches using mutants with arrayed DNA content < 24**

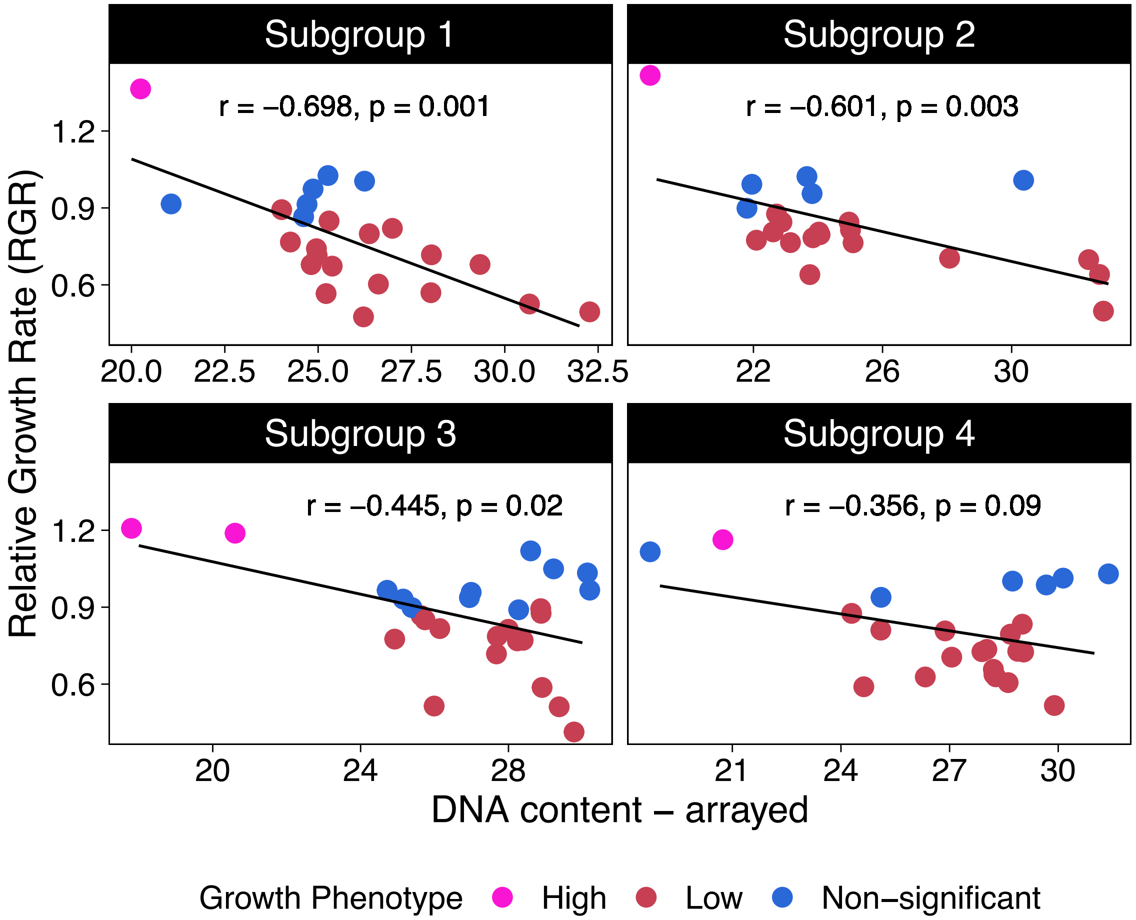


**Supplementary Figure 15. Relationship between growth rates and replication potential.**

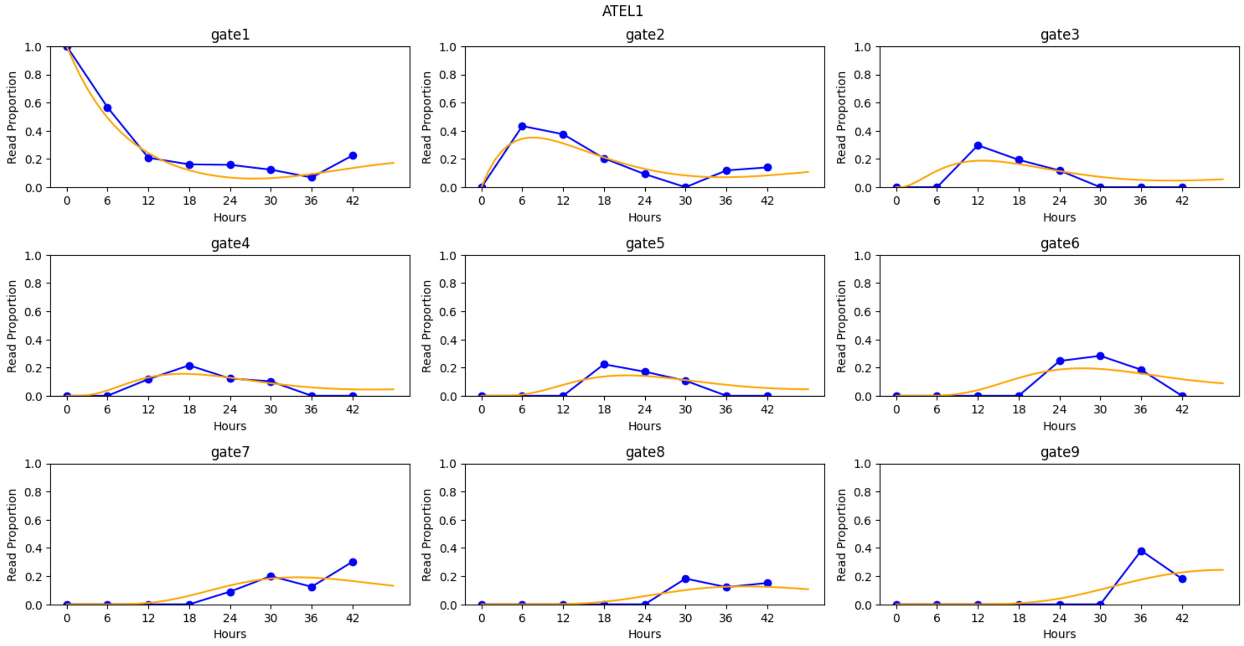
ATEL1
gate1
gate2
gate3
gate4
gate5
gate6
gate7
gate8
gate9
Read Proportion
Hours
0 6 12 18 24 30 36 42
0.0 0.2 0.4 0.6 0.8 1.0

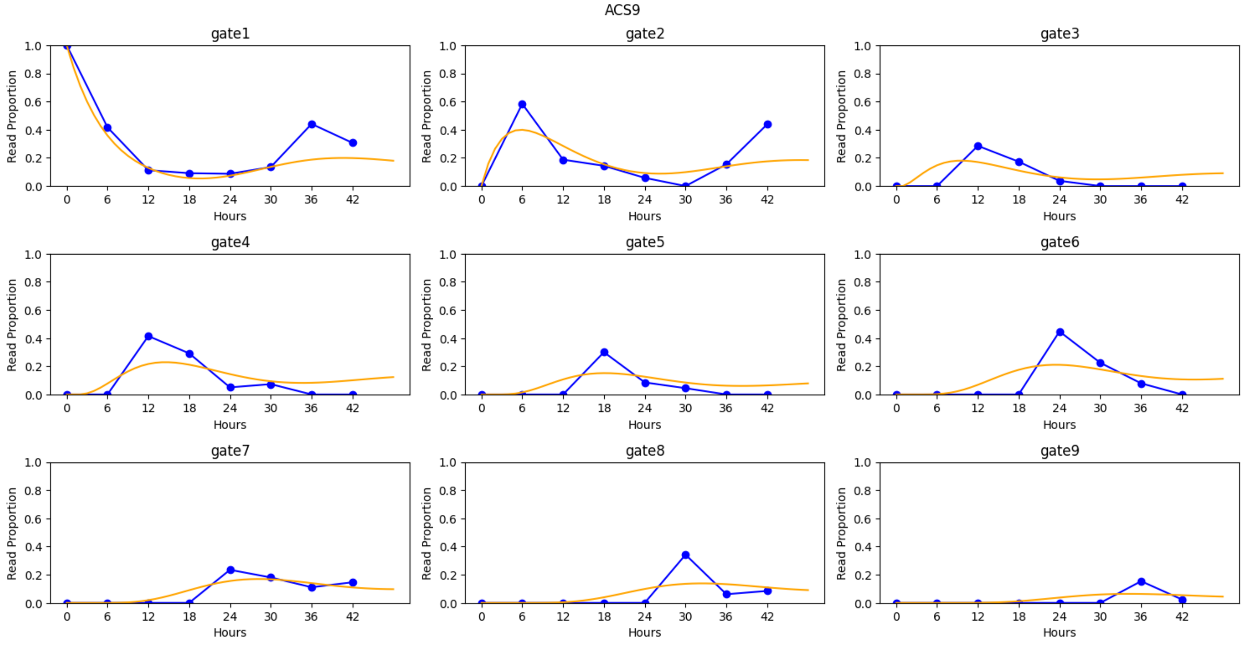
ACS9
gate1
gate2
gate3
gate4
gate5
gate6
gate7
gate8
gate9
Read Proportion
Hours
0 6 12 18 24 30 36 42
0.0 0.2 0.4 0.6 0.8 1.0

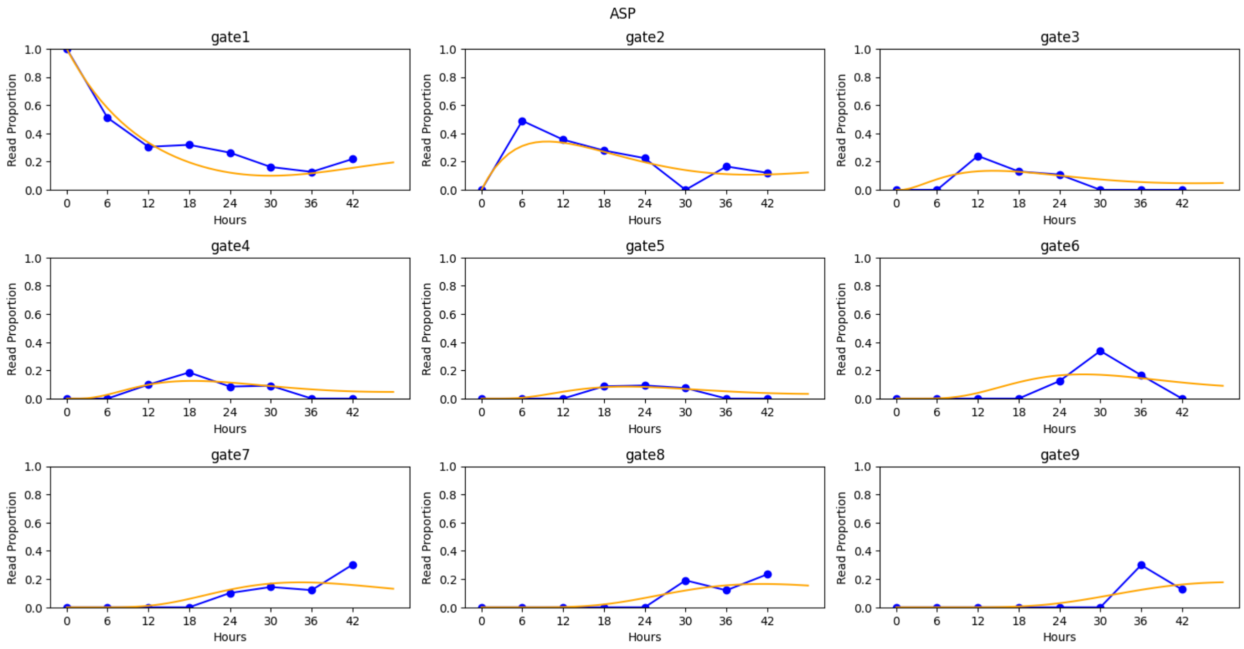
ASP
gate1
gate2
gate3
gate4
gate5
gate6
gate7
gate8
gate9
Read Proportion
Hours
0.0
0.2
0.4
0.6
0.8
1.0
0
6
12
18
24
30
36
42

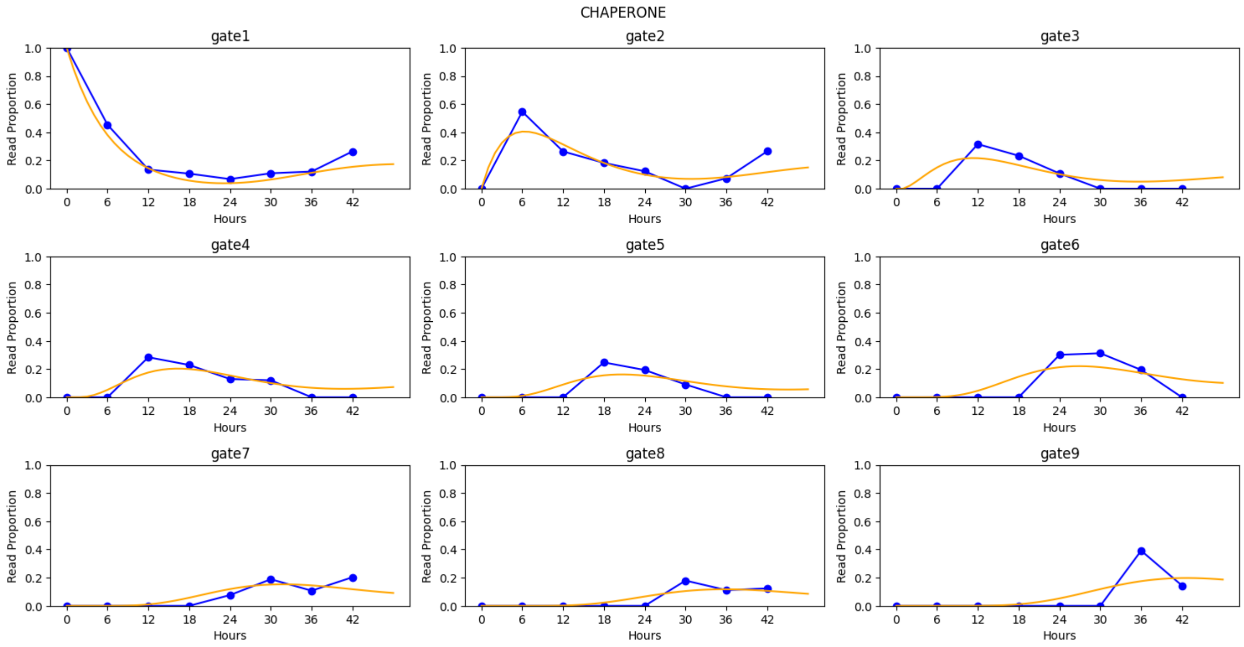
CHAPERONE
gate1
gate2
gate3
gate4
gate5
gate6
gate7
gate8
gate9
Read Proportion
Hours
0.0
0.2
0.4
0.6
0.8
1.0
0
6
12
18
24
30
36
42

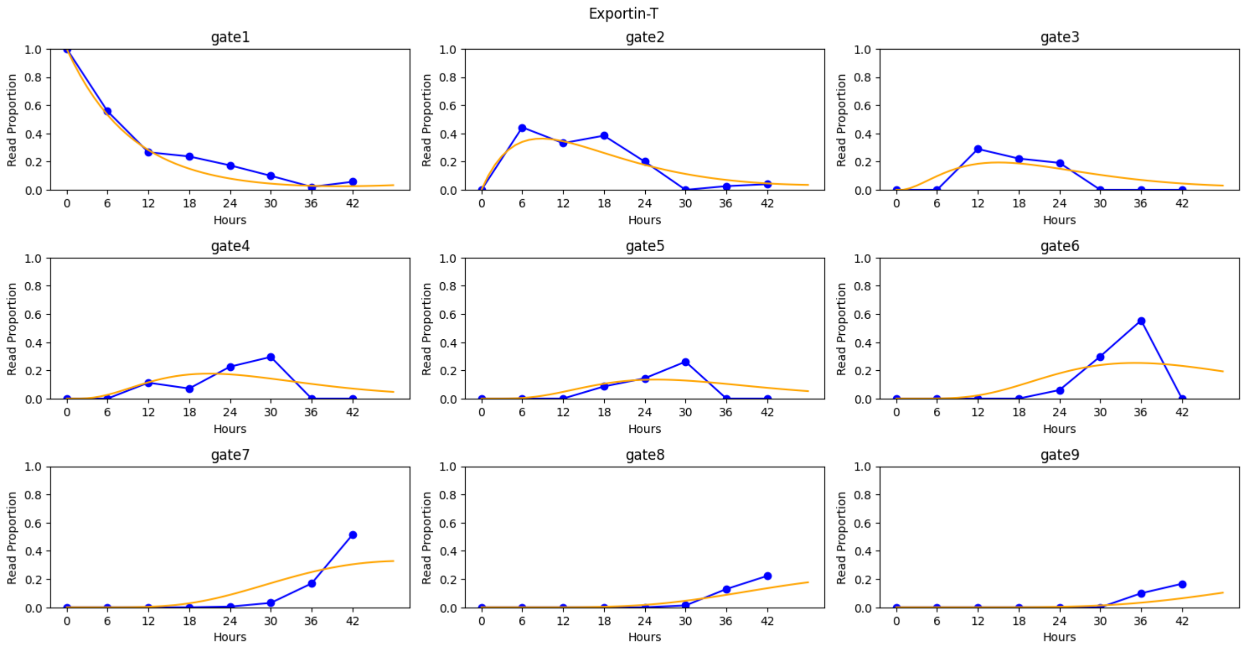


**Supplementary Figure 16**. Simulated (orange) and observed (blue) read proportions in each gate for genes in the proof of principle experiment.

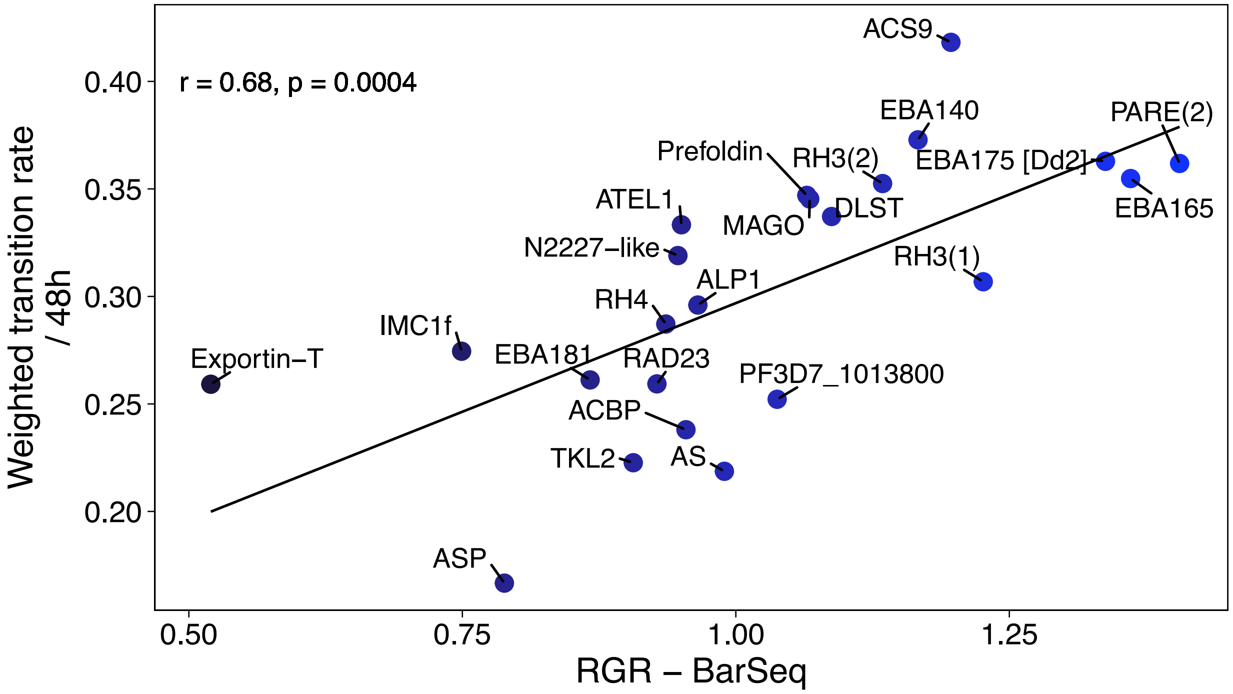


**Supplementary Figure 17**. Correlation between rates of transition of deletion in the IDC with RGR estimated using BarSeq.

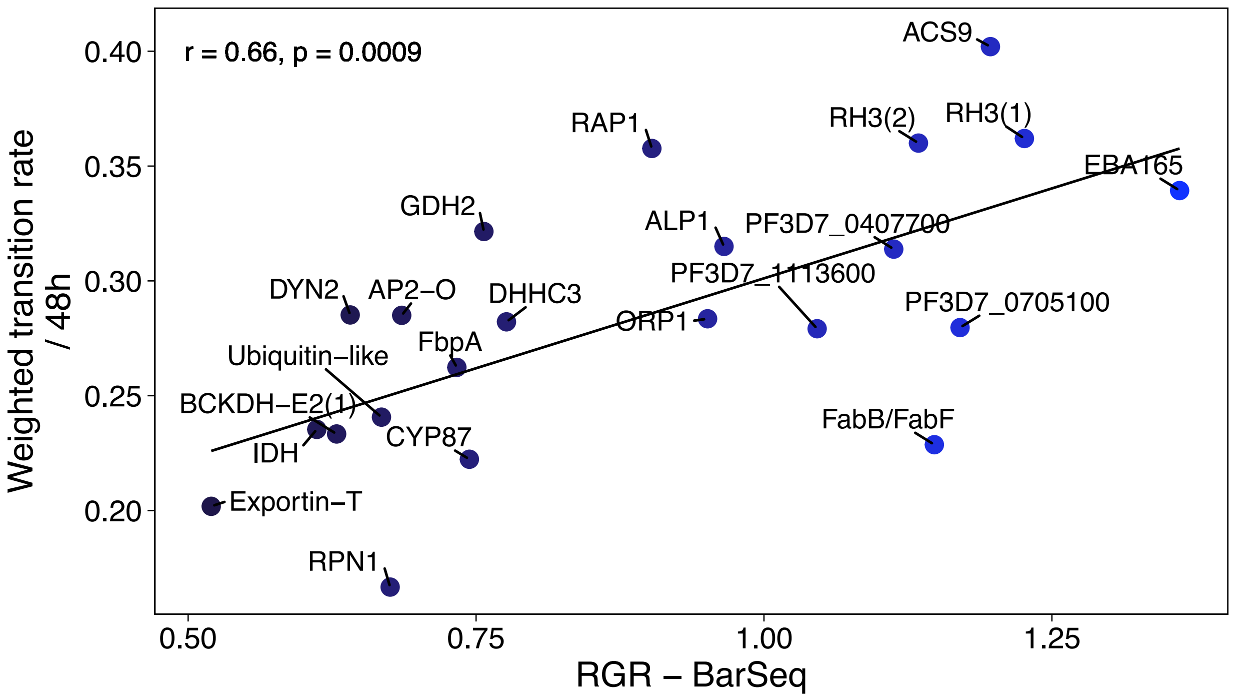


**Supplementary Figure 18**. Correlation between rates of transition of deletion in the IDC with RGR estimated using BarSeq.

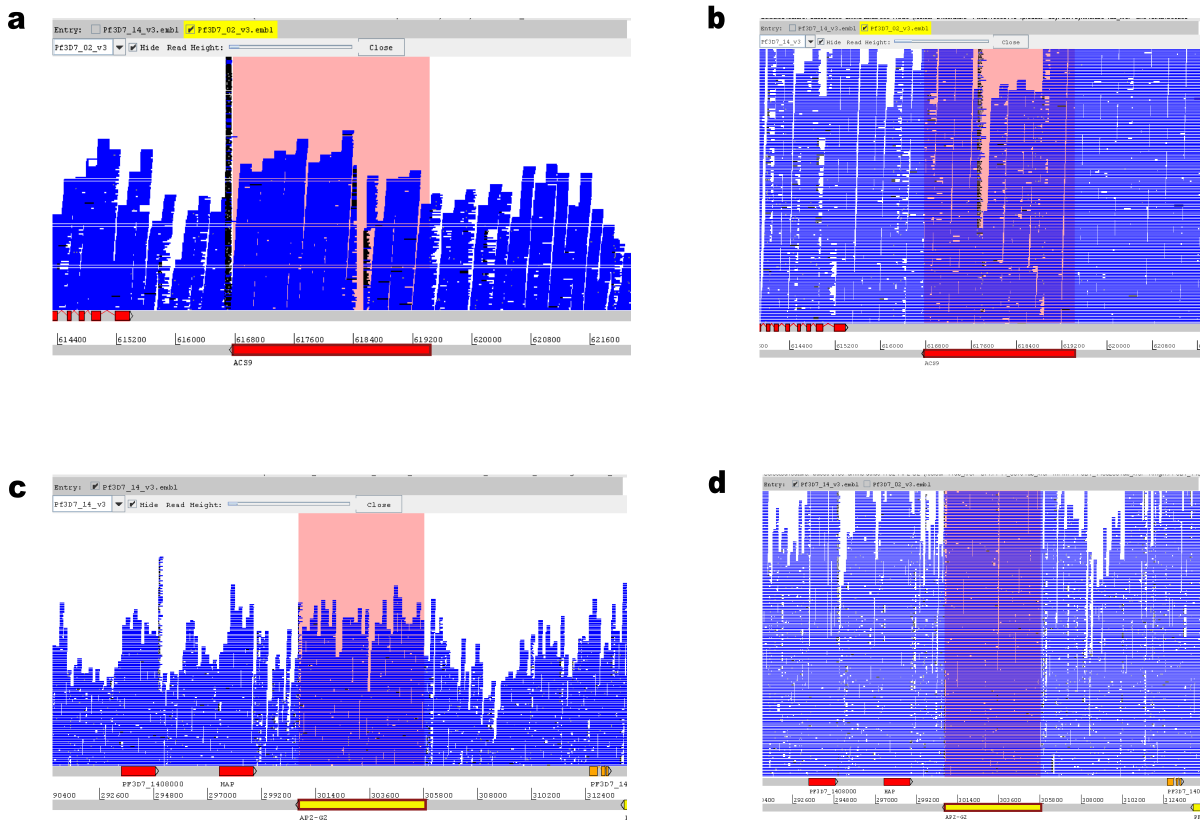


**Supplementary Figure 19**. Visualization of whole genome sequence reads using Artemis genome browser. (a-b) – ACS9 deletion line, (c-d) - NF54 wild type.
